# Connectivity Map and perturbation-based sensitivity analysis identifies MEK inhibitors as senolytics in human lung fibroblasts

**DOI:** 10.1101/2024.11.16.623933

**Authors:** Amirhossein Nayeri Rad, Simon Sperger, Leigh M. Marsh, Konrad Hoetzenecker, Ingo Lämmermann, Johannes Grillari

## Abstract

Recently, the elimination of the disease-associated accumulation of senescent cells using senolytics has been shown to exert health benefits in animal studies. However, due to the heterogeneity of cell senescence and its unrecognized master regulators, drug development faces a complexity that must be handled. Bioinformatic elucidation of genes and pathways involved in senolysis and prediction of senolytic activity of compounds can cut costs and facilitate faster achievements in the field. In the present investigation, after obtaining the consensus gene signature of senescent fibroblasts of lung origin and deriving its anti-apoptotic module, we utilized Connectivity Map (CMap) alongside small molecule and genetic perturbation sensitivity data in cancer cell lines to identify drugs and genetic interventions that might induce apoptosis or sensitize senescent cells to apoptosis. Through bioinformatic evaluations, we speculate that activation of early stages of autophagy which contributes to the formation of autophagosomes, concurrent with the activation of waste protein concealment system by the mean of p62 and chaperoning system alongside an increase in JUNB gene expression can secure the survival of the senescent cells even when homeostasis of different cellular processes is disrupted. Moreover, our bioinformatic evaluation proposed selumetinib, a MEK inhibitor, as a senolytic against senescent lung fibroblasts. The senolytic activity of a variety of MEK inhibitors in senescent lung fibroblasts was confirmed using human lung fibroblasts in vitro.

## Introduction

Nowadays cancer, aging, and many hyper-inflammatory diseases have been revealed to be linked to cellular senescence [1]. Thereby, understanding senescence and its underlying signaling pathways might become one of the game-changers in the therapy of a vast range of diseases termed ‘senopathies’ [2], in which senescent cells are causally involved in the pathogenesis. It has been indicated that the genetics of senescence, depending on the cell origin, is heterogenous and involves different elements [3, 4]. This heterogeneity has also contributed to a hurdle in universal therapeutic targeting of these cells. A class of these drugs, called senolytics, which induce apoptosis in senescent cells, e.g., dasatinib+quercetin [5], fisetin [6], navitoclax [7], and others [8] have been shown to exert their effect in a cell-type specific manner. In a previous study, we integrated a set of bioinformatic measurements including transcriptional connectivity attained from Connectivity Map (CMap) and network proximity scores to discover another class of senotherapeutics [4], called senomorphics, which attenuate the detrimental functions of senescent cells like their inflammatory secretions, named senescence-associated secretory phenotype (SASP) [9].

CMap is a large-scale compendium of the cellular effects of small-molecule and genetic perturbations that calculates the connectivity of the perturbations’ effect with the signature which can be requested through the CLUE Query online platform [10]. While CMap data are abundantly related to cancer cell lines (not normal cells), a variety of studies have used CMap negative connections to repurpose drugs for a wider range of diseases rather than cancer, like aging [11], Alzheimer’s disease [12], osteoporosis [13], COVID-19 [14], and bipolar disorder [15], and some studies have utilized CMap positive connections to identify mimetics inducing a specific phenotype e.g. new caloric restriction mimetics [16]. Although many studies have taken advantage of CMap in drug discovery, due to the multiplicity of factors (including cell type, duration, dose, etc.) it is still a challenge to increase the predictive performance of the methodologies used for ranking the interventions [17].

In this study, we aimed to discover new senolytics and senolytic targets and gain insight into the underlying mechanism of apoptosis resistance in senescent cells using bioinformatic methodologies and publicly available data. To overcome the heterogeneity problem of different senescent cell types we focused on senescent fibroblasts of lung origin which had numerous RNA sequencing datasets availably deposited in the Gene Expression Omnibus (GEO) database. In addition, administration of senolytics in clinical study of idiopathic lung fibrosis (IPF) has shown to be safe and to at least increase gait speed and 6-min walk distance (6MWD) of IPF patients [18]. IPF was chosen for the clinical trial due to a clear increase of senescent cells in pre-clinical mouse models as well as in human tissue samples [19]. However, also other lung diseases qualify as senopathies, including chronic obstructive pulmonary disease (COPD) and lung aging [20].

Therefore, we here report that by compilation of differential gene expression datasets, we acquired a consensus senescence signature of human lung fibroblasts. We then performed a massive interpretation of this senescence signature using bioinformatic gene-set enrichment analysis coupled with a gene-by-gene literature review. Then we extracted the anti-apoptotic module of the signature and scrutinized its association with the pool of transcriptional effects of small-molecule and genetic perturbations attained by CMap. As senolysis is mainly driven by induction of cell death and apoptosis in senescent cells, we further consolidated our CMap-based prediction of senolytic activity with the real large-scale cell viability assessments of the perturbed cells, obtained from the cancer dependency map (DepMap) portal. DepMap which is an online platform for systematic identification of genetic dependencies and small molecule sensitivities [21], has provided the datasets of genetic loss of function sensitivities from the project Achilles [21] and small molecule sensitivities from The Cancer Therapeutics Response Portal v2 (CTRP v2) [22]. Finally, we tested one resulting and promising candidate class, MEK inhibitors, for their potential activity as senolytics using human lung fibroblasts (HLFs).

## Results

### Unraveling the consensus senescence signature

Differential gene expression was analyzed in 14 datasets from 10 different studies of senescence induction using 3 different senescence inducers (Ras-oncogene induced - OIS, irradiation-induced IRIS, and replicative senescence - RS) versus proliferating control cells in 2 fibroblast cell lines of lung origin, WI-38 and IMR-90 (Dataset S1. Sheet1). We performed principal component analysis (PCA) to see how the main principal components of these differential gene expression profiles distribute against each other in a simplified two-dimensional space (Fig. 2A). Differential gene expression profiles in consequence to the 3 different senescence inducers show low inter-cellular variations, while variations due to different inducers were larger. This makes senescent cells of the same origin classifiable based on their mode of induction. Still, we speculated that several of the gene expression levels of these cells including cell cycle arrest, resistance against apoptosis, or their secretory profile must be independent of the senescence inducer. Therefore, we calculated a consensus signature independent of the senescence inducer including 384 upregulated genes and 605 downregulated genes (overall 989 genes) (Dataset S1. Sheet2). An overview of the bioinformatic workflow is shown in Fig. 1.

**Fig. 1:**
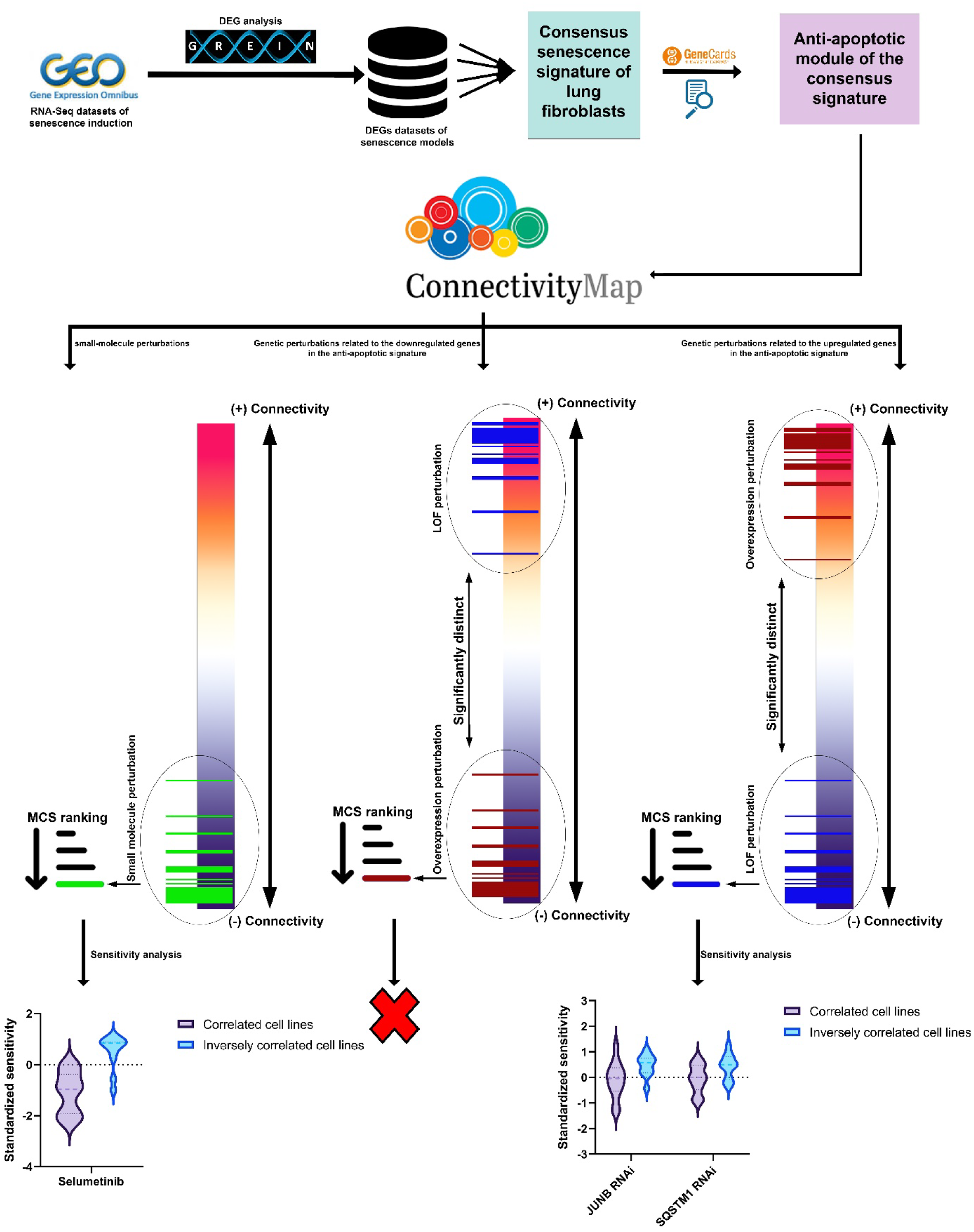
Schematic of the bioinformatic pipeline of the study.

**Fig. 2:**
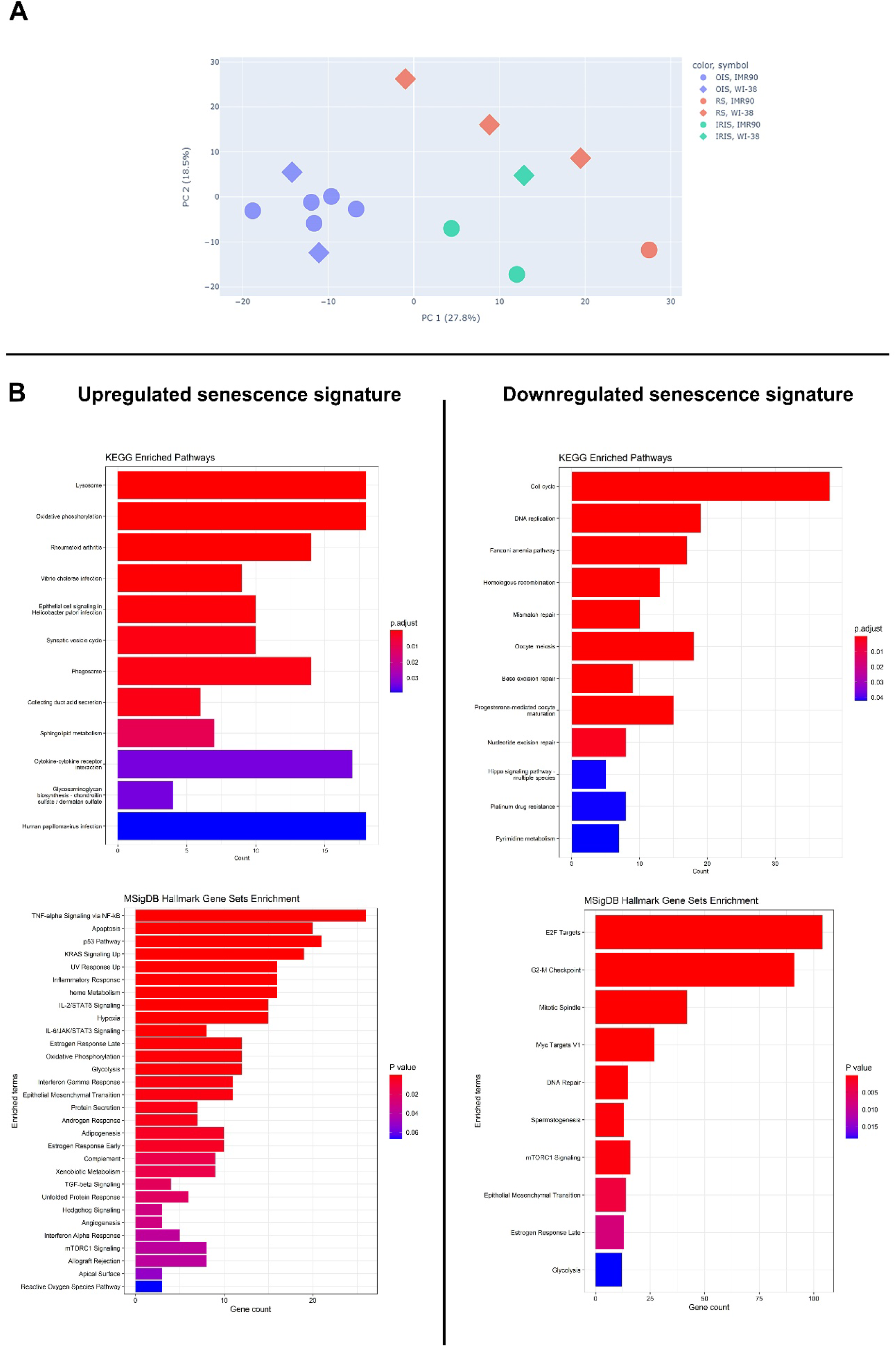
Principal component analysis of the differential gene expression profiles of the collected senescent models, and enrichment analysis of the consensus signature derived from all these datasets. (A) The first two principal components of differential gene expression profiles, are labeled by the senescence induction model and the cell type. (B) KEGG pathways and MSigDB Hallmark gene set enrichment analysis for up and downregulated signature separately. OIS: oncogene-induced senescence, RS: replicative senescence, and IRIS: irradiation-induced senescence.

CDKN1A (p21), one of the prominent markers of senescence, that enforces cell cycle arrest, was included in the signature of upregulated genes of HLFs, while surprisingly, CDKN2A (p16) or TP53 (p53) two other well-known genes involved in the regulation of the cell cycle in senescence were not included. By examining each differential gene expression dataset individually for a consensus senescence signature, we found that p16 was unanimously upregulated, and p53 unexpectedly was downregulated in all OIS datasets. However, this unanimity in either upregulation or downregulation was not observed in RS or IRIS datasets. These emphasize p21 as a better transcriptional diagnostic marker in senescent lung fibroblasts. It must be taken into account that we are just noting the transcription quantity; the protein levels might be determined by post-transcriptional control, too. Moreover, alternative splicing/reading of CDKN2A gene might have affected its undelineated transcriptional level. MSigDB Hallmarks 2020 enrichment analysis of the consensus signature by the Enrichr R tool demonstrated that the upregulated genes were significantly overrepresented in “TNF-alpha Signaling via NF-κB”, “p53 Pathway”, and “KRAS Signaling Up” (Fig 2. B). All three pathways are well connected to senescence: activation of the NF-κB pathway not only induces the SASP, but is also implicated in senescent cell survival [23], p53 signaling plays a role in directing the fate of cells towards senescence or apoptosis upon cellular stress [24, 25], while upregulation of RAS in senescence can ensure cellular survival or lead to developing a senescence-associated phenotype by two distinct but interconnected pathways the RAF/MEK/ERK signaling cascade and the PI3K/AKT pathway [26, 27].

KEGG pathways enrichment analysis of the consensus underexpressed signature, represents the deactivation of pathways related to “Cell cycle”, “DNA replication”, “Fanconi Anemia pathway” and DNA repair mechanisms including “Homologous recombination”, “Mismatch repair”, “Base excision repair”, and “Nucleotide excision repair”. Gene ontology (GO) overrepresentation analysis of the downregulated signature beyond the processes involved in cell cycle, DNA repair, and replication manifests downregulation of the genes which control the epigenome and packaging of the DNA (Fig. S1 and Dataset S1. Sheet3).

### Manually curated senescence anti-apoptotic signature and its connectivity map

The consensus senescence signature was then reduced to only the anti-apoptotic module of the whole signature (anti-apoptotic signature, ASS). On a cautious note, it needs to be taken into account that the attained anti-apoptotic signature does not represent all the genes involved in resistance to cell death or apoptosis, since the method we utilized was designed in a way to decrease the effort needed to manually select (through vast literature mining) the genes with potential anti-apoptotic role in senescence and to reduce the risk of error in our selections. This became true by initially sorting the genes related to apoptosis using Genecards’ algorithmic gene scoring system which helped us to omit the genes that did not have enough evidence in the scientific literature to support their inclusion. However, in order not to miss important genes related to cell survival, prior to the manual literature-mining step, we appended the genes which were annotated by GO to be involved in regulation of cell death, to the list of the genes. Still, there is the possibility that some of the genes which were not well discussed in regard to apoptosis in the literature have been missed. Ultimately, we defined a signature including 59 upregulated genes and 57 downregulated genes whose inverse regulation have been shown to be potentially fatal or sensitize cells to apoptosis (table. 1). The evidence for and against the inclusion decision of the selected genes are provided as supplementary document (Supplementary file 1). A transcriptional connectivity map of the anti-apoptotic signature was attained by entering up and downregulated signature into the CLUE Query platform.

**Table. 1:**
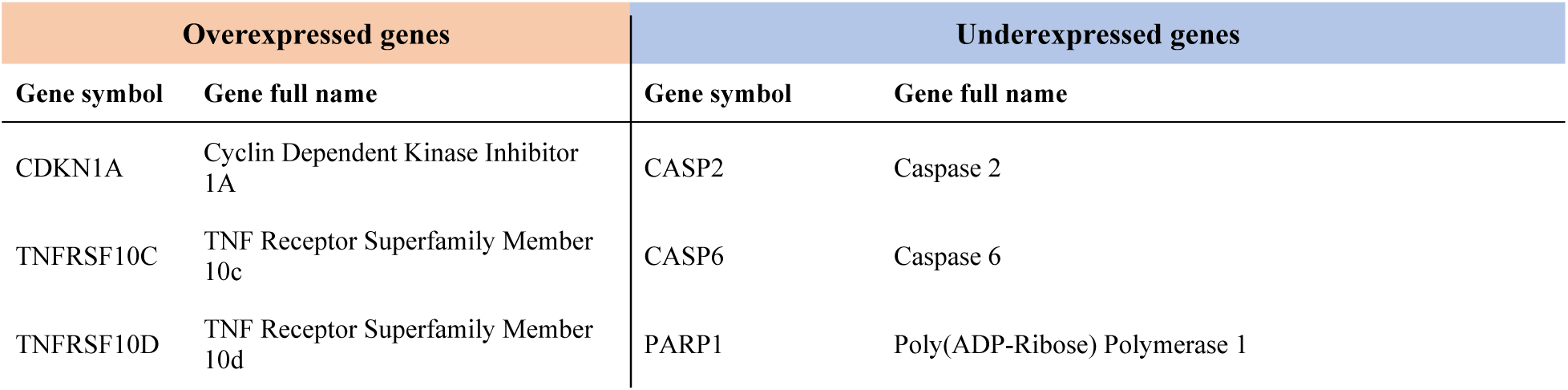
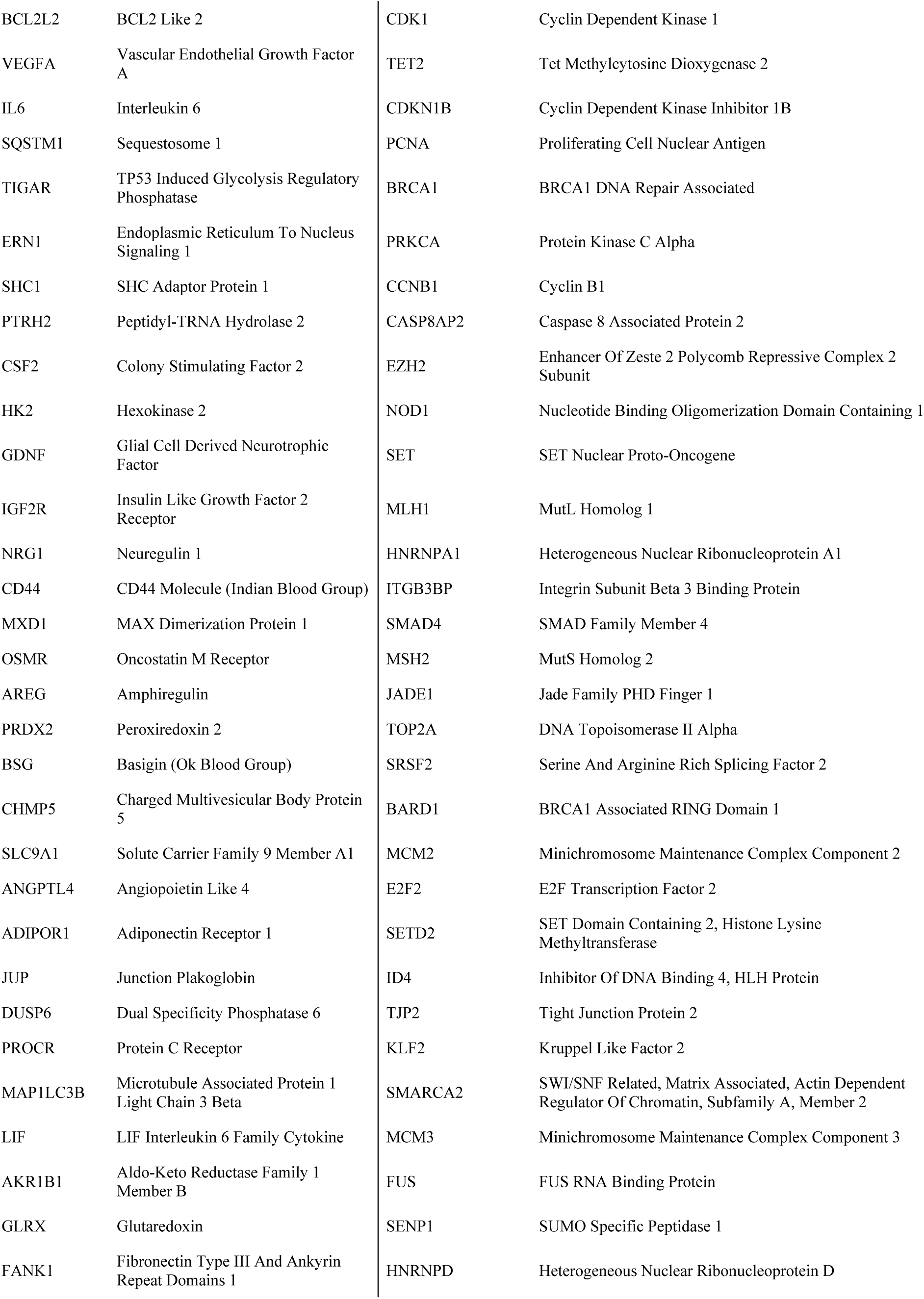
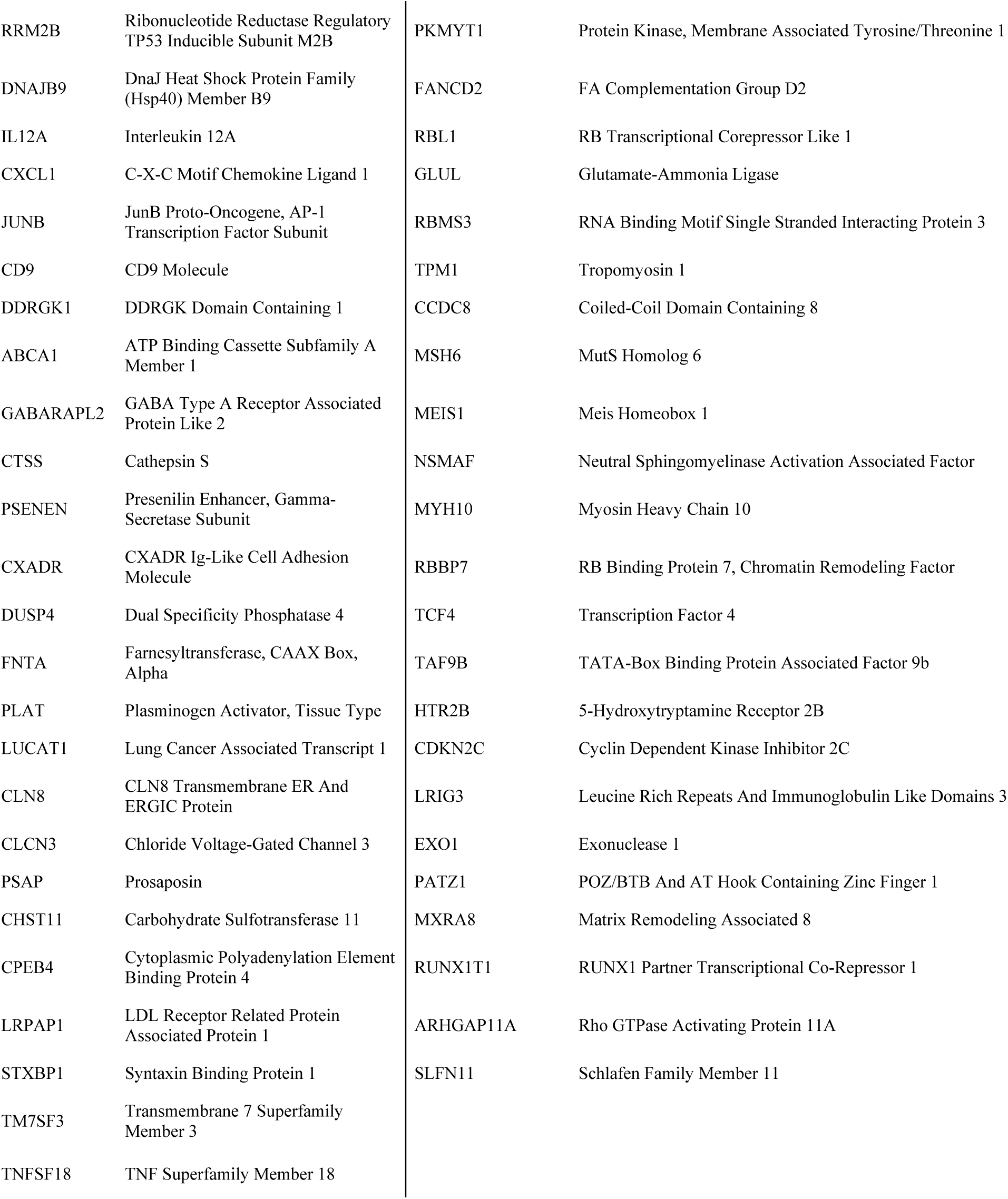
The anti-apoptotic module of the senescence signature (anti-apoptotic signature).

### Discovering potential senolytics utilizing CMap and compound sensitivity assessment

CMap retrieved the connectivity of 136460 small-molecule perturbations. Excluding perturbations in negatively correlated cell lines, 116128 perturbations relating to 21858 specific small molecules (compounds) were reached. To perform our analysis, we eliminated data of small molecules which had less than 10 perturbations each and their median TAS was less than 0.1 (TAS has been proposed as a factor representing the transcriptional strength and inter-sample correlation of each perturbation [10]) which left perturbational data related to 3628 unique compounds (Dataset S2. Sheet1-4). In order to sort small molecules over their association with the anti-apoptotic signature, we calculated the mean of normalized connectivity scores (NCSs) of each small-molecule entity over its different perturbations (in different cell types, or used concentration, even some were fully replicated in different institutes) and the percentile of this mean connectivity score (MCS) among the scores of all 3628 unique compounds. Previously we used mean squared distance (MSD) scoring to increase the cost of outliers in the final ranking of compounds [4]. However, this time we suspected that some of the NCS outliers appeared by the extremely different genetic context in which perturbations were exerted. Thus, we removed the perturbations in cell types whose gene signatures were negatively correlated with the anti-apoptotic signature (these cells were: ‘HL60’, ‘JURKAT’, ‘KMS34’, ‘MINO’, ‘NALM6’, ‘NOMO1’, ‘OCILY10’, ‘OCILY3’, ‘SHSY5Y’, ‘THP1’, ‘U2OS’ and ‘U937’) and we did not include their NCSs for calculating the MCS. Thus, we included the remaining data equally (even if any outliers remained after neutralizing the context dependency of the scores) and we further did not think that we needed to penalize the NCSs that were far from the expected result. The MCSs were further used for ranking the compounds.

The gene signature, which was used as the representative of each cell line was taken from the standardized gene expression matrix of the Cancer Cell Line Encyclopedia (CCLE) gene expression profiles obtained from the Harmonizome database. The positive MCS of a compound represents direct connectivity of its transcriptional impact with the signature of interest, which would implicate a potential senescence inductive effect. However, the negative MCS of a compound represents inverse connectivity of its transcriptional impact with the signature of interest, which we expect to exert a pro-apoptotic effect on the corresponding cells.

The top 10 percentile of the MCSs, with the lowest score of 0.68, included many of the compounds previously indicated to be senescence inducers e.g., etoposide, palbociclib (which is a CDK4/6 inhibitor), vincristine, topotecan, bortezomib, camptothecin, and vinblastine [28] (Dataset S2. Sheet6). Carfilzomib, which yet has not been studied for its association with senescence emergence, and bortezomib both are proteasome inhibitors that we previously mentioned to force-overexpress the genes involved in autophagosome formation, mimicking the anti-apoptotic senescence phenotype. Bortezomib and carfilzomib with MCSs of 0.99 and 1.46, placed at the 96.52 and 99.99 percentile of the global MCSs, respectively. It must be remarked that doxorubicin which is widely used as a senescence inducer got an MCS of 0.61 and posed at the 87.95 percentile of MCSs. Intriguingly, thapsigargin which we speculated to simulate senescence transcriptional signaling through endoplasmic calcium regulation, has got an MCS of 0.8 which is placed at the 92.88 percentile of MCSs pool. Since there was not a sufficient number of compounds that have CMap profiles and have been experimentally confirmed to have senolytic effects in lung fibroblasts (even some like fisetin have failed to show senolytic activity against senescent lung fibroblasts [29]), we were not able to validate in silico whether having an inverse connection has any association with senolytic effects. However, by assuming that the least connected compounds might potentially be senolytic, we selected the bottom 10 percentile of the compounds (MCSs were in the range of −1.07 to −0.52) for a more detailed analysis in the rest of the study (Dataset S2. Sheet7).

Among the 363 selected compounds which were inversely connected to the anti-apoptotic signature, we sought to see if any of these compounds would be known to selectively kill cancer cell lines of similar signatures (directly correlated with both the whole consensus senescence signature and the antiapoptotic signature) against cell lines with dissimilar signature (inversely correlated with both the whole consensus senescence signature and the antiapoptotic signature). Therefore, the standardized CCLE gene expression data were used for identifying the most and the least correlated cell lines’ signatures. We used the standardized CTD^2^ drug sensitivity dataset for comparing the compound lethality in correlated cell lines vs inversely correlated ones. The cell lines should have been shared between the CCLE dataset and the CTD^2^ sensitivity dataset. In addition, to avoid bias towards any specific lineage of cancer cell lines, at most the two most correlated cell lines in each lineage were included in the analysis). Selected cell lines with respect to their lineage are represented in the table. 2.

**Table. 2.**
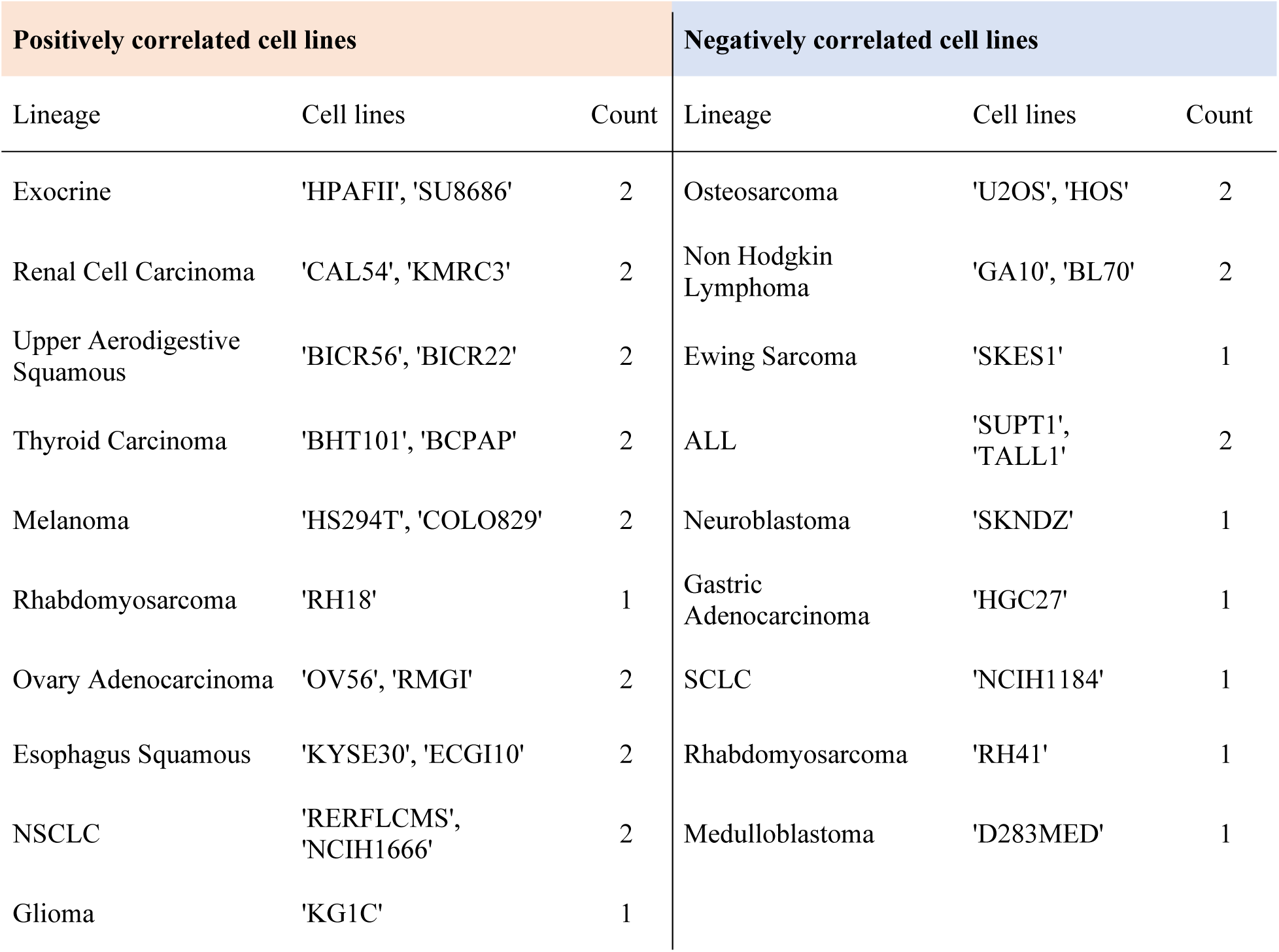
The selected positively correlated cell lines and the inversely correlated cell lines, with respect to their cancer lineages, which were shared between CCLE and CTD^2^ datasets and were utilized in the comparative assessment of small molecule sensitivities.

Sensitivity data of merely 11 out of the 363 previously selected compounds were available from CTD^2^ dataset (Dataset S2. Sheet8). Of these 11 compounds, 2 compounds – selumetinib which is a MEK inhibitor, and SJ-172550 which is an MDMX inhibitor – were demonstrated to exert higher toxicity in correlated cell lines vs inversely correlated ones with p-values of 0.00002 and 0.0046, respectively, whereby significance was analyzed by permutation test.

To better gain insight into the compound sensitivities in the selected cell lines with respect to all other CTD^2^ cell lines, we calculated the percentile of the mean sensitivity in correlated cell lines and inversely correlated cell lines separately. Data is reported in table. 3. Next, we evaluated replicate level viability data (replicate-level viability dataset which is provided by DepMap, includes the detailed measured viabilities of cells in exposure to different concentrations of the compounds, each in replicates, which are all reported) of selumetinib and SJ-172550 to ensure that the effects of the compounds on cell numbers in correlated cell lines vs inversely correlated cell lines is indeed due to toxicity and would not be confounded by a potential proliferation-enhancing effect in inversely correlated cell lines (Dataset S2. Sheet9). The mean viability of both correlated and inversely correlated cell lines exposed to SJ-172550 were more than 1 which was indicative of a proliferation-enhancing effect mostly toward inversely correlated cell lines. Therefore, through our bioinformatics assessments, selumetinib remained the only compound that we identified as a candidate senolytic for experimental testing.

**Table. 3.**
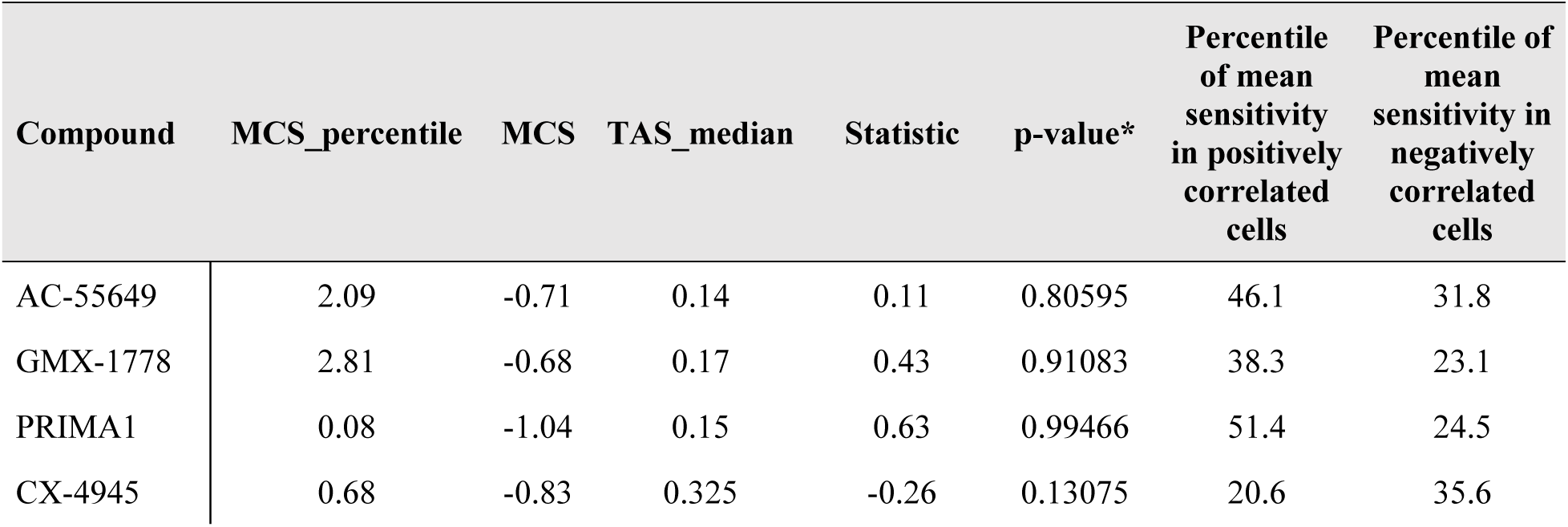
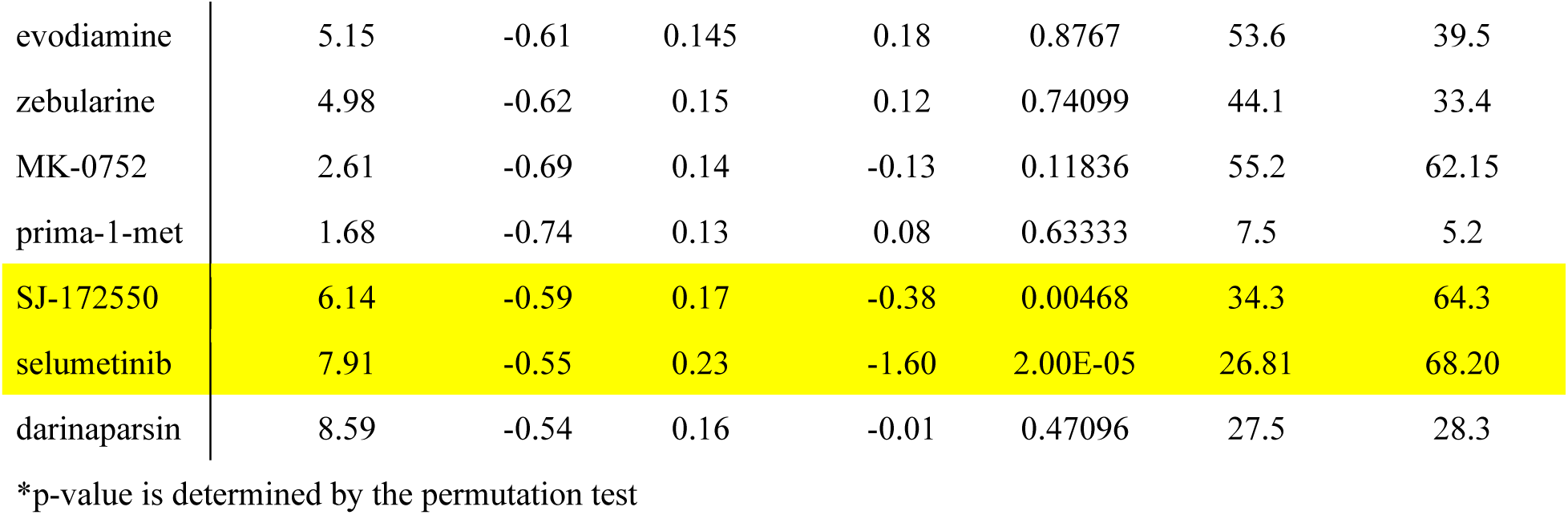
Analysis of sensitivity to the shared bottom 10 percent connected compounds. The Mean Connectivity Score (MCS) and its percentile (MCS_percentile) alongside the median of TAS (TAS_median) are derived from Connectivity Map. The difference between the compound sensitivity in the correlated cell lines and the inversely correlated ones is defined by “Statistic” and “p-value”. Moreover, the percentile of mean sensitivity in each group is represented.

### Bioinformatic uncovering of the gene regulatory system underlying senescence resistance to apoptosis

We utilized similar methodologies as we used for bioinformatic analysis of small molecules with some modifications, to find out the underlying signaling which might contribute to the senescence apoptosis resistance. By gaining knowledge about the master regulators of this cellular anti-apoptotic trait, researchers would be able to target these genes or processes to discover high efficiency senolytics. Genetic perturbations in the CMap, contrary to the small-molecule perturbations, are bi-directional, which means that the direction of the regulation of the gene had to be taken into account in our analyses. We used two different approaches to evaluate the connectivity of single gene regulation with the emergence of the anti-apoptotic signature (schematically represented in Fig. 1). First, we applied the same MCS-based ranking for the perturbations that negatively regulate the gene direction of expression in the anti-apoptotic signature. It means that for the overexpressed genes in the anti-apoptotic signature we employed the NCSs of the loss of function perturbations (including CRISPR and shRNA knockdown) and for the underexpressed genes we used the NCSs of the gain of function perturbations (overexpression). The more negative MCS would hypothetically demonstrate that the inverse regulation of the corresponding gene can potentially counteract the whole anti-apoptotic signature, leading to cell death. Genes with the 5 smallest MCSs are represented based on 3 different functions of their perturbations (overexpression, CRISPR knockdown, and shRNA knockdown) (Dataset S3. Sheet2-4). We also pooled the data of both CRISPR and shRNA knockdown perturbations and calculated the MCSs based on the NCSs of the whole pooled loss of function perturbations (gene interventions with the 10 least MCSs are represented in the Dataset S3. Sheet5). Second, regardless of the ranking of the genes, we looked if the regulation of each gene in two opposite directions could differentially drive the cells towards two significantly distinct transcriptional fates: the one with the same direction as the direction of the gene in the anti-apoptotic signature, towards senescence induction (more specifically, towards raising the senescence apoptosis-resistance machinery), and the other one against the direction of the gene in the anti-apoptotic signature, towards undermining the pro-survival signalings. To carry it out, for the genes in the overexpressed anti-apoptotic signature we examined if the MCS of their overexpression is significantly larger than the MCS of their pooled loss of function, and for the genes in the underexpressed anti-apoptotic signature we examined if the MCS of their pooled loss of function is significantly greater than the MCS of the overexpression (significance was evaluated by permutation test).

The results of this analysis are shown in the Dataset S3. Sheet6-7. Collectively, based on the rankings and the impact of single genes in the active shaping of the anti-apoptotic vs the alleged apoptotic signature, we identified potential key players of this system. According to our data, we suggest that EZH2 and PCNA are the two most determining genes in the underexpressed anti-apoptotic signature. Among the upregulated genes, JUNB, SQSTM1, PRDX2, and CDKN1A (p21) are the master transcriptional regulators whose impairment of each might presumably be a powerful tool to induce apoptosis in senescent cells (especially lung fibroblasts). In the next step, similar to the bioinformatic test that we designed for analyzing cell death upon exposure of the senescence-mimicking cell lines to small molecules, due to the availability of gene LOF sensitivity data in cell lines (sensitivity data related to gene gain of function were not available), we could inspect through real sensitivity data to identify that the depletion of which overexpressed genes would be potentially fatal in senescent fibroblasts. Correlated cell lines and inversely correlated cell lines which were used for comparative sensitivity analysis of the CRISPR and RNAi knockdowns are shown in Dataset S1. Sheet5. Our bioinformatic assessments demonstrate that JUNB loss by either CRISPR or RNAi depletion led to a significantly increased cell death in correlated cell lines in comparison to the inversely correlated ones (Dataset S3. Sheet8-9).

In addition, RNAi LOF perturbation hints towards the expression of SQSTM1, LIF, and AREG as important genes in counteracting apoptosis in cell lines mimicking the here identified senescence signature and its anti-apoptotic module (Table. 4).

**Table. 4.**
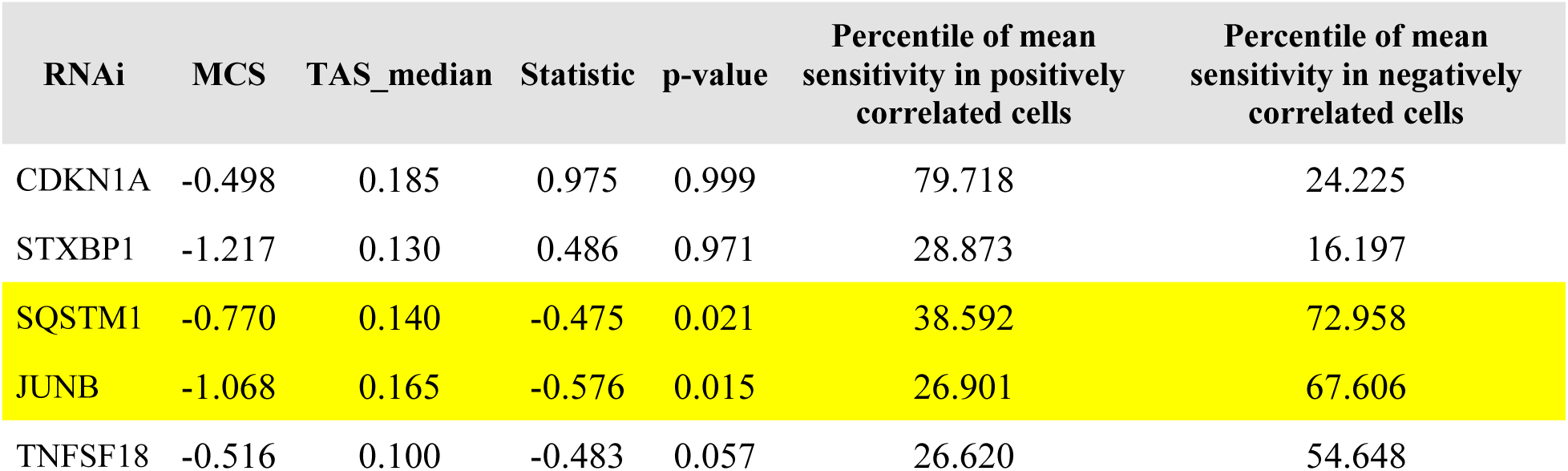
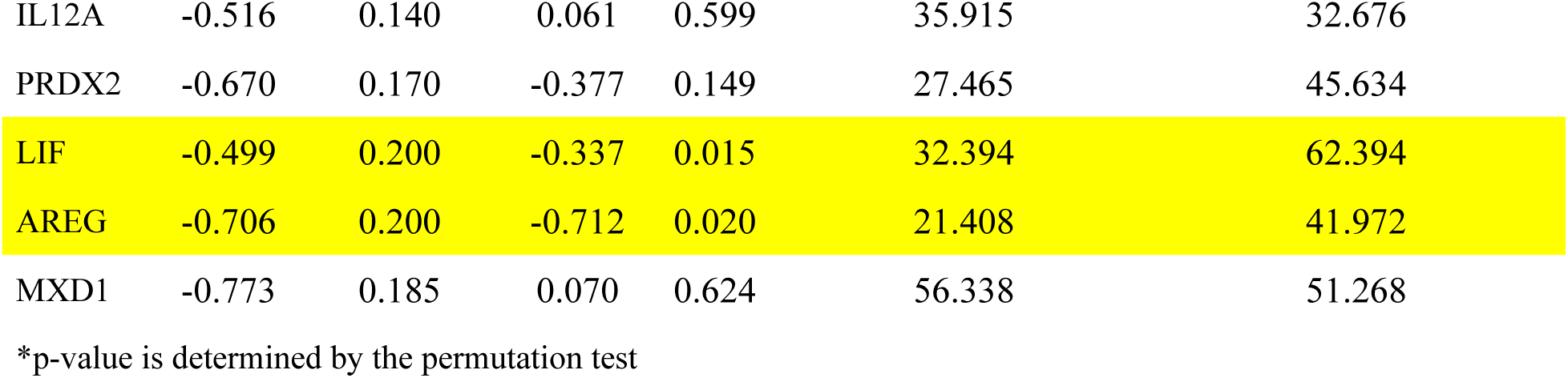
Analysis of sensitivity to the RNAi loss of function (LOF) perturbations of the genes whose LOF were among the top 10 inversely connected interventions with the anti-apoptotic signature. The Mean Connectivity Score (MCS) and the median of TAS (TAS_median) are derived from Connectivity Map. The difference in the RNAi sensitivity in correlated cell lines vs inversely correlated ones is defined by “Statistic” and “p-value”. Moreover, the percentile of mean sensitivity in each group is represented.

### MEK inhibitors are senolytic in human lung fibroblasts in vitro

To confirm the bioinformatic prediction regarding the senolytic activity of selumetinib in senescent lung fibroblasts, quiescent versus senescent human lung fibroblasts, HLF102, were treated with selumetinib. Stress-induced premature senescence (SIPS) was triggered using doxorubicin as shown in the scheme (Fig. 3A) as well established in previous works [30–32]. Interestingly, selumetinib selectively killed senescent cells with an IC50 of around 8.35 µM, while quiescent control IC50 values were at 51.82 µM (Fig. 3B). To test, if the senolytic activity can be generalized to the whole class of MEK inhibitors, we selected four additional MEK inhibitors: Binimetinib, Cobimetinib, Mirdametinib and Tramentinib. Indeed, all here tested MEK inhibitors displayed senolytic activity (Fig. 3C-F), with Trametinib showing an IC50 in the nanomolar range with 0.06 µM and a fold difference to quiescent control cells of around 70 fold with an IC50 of 4.14 µM.

**Fig. 3:**
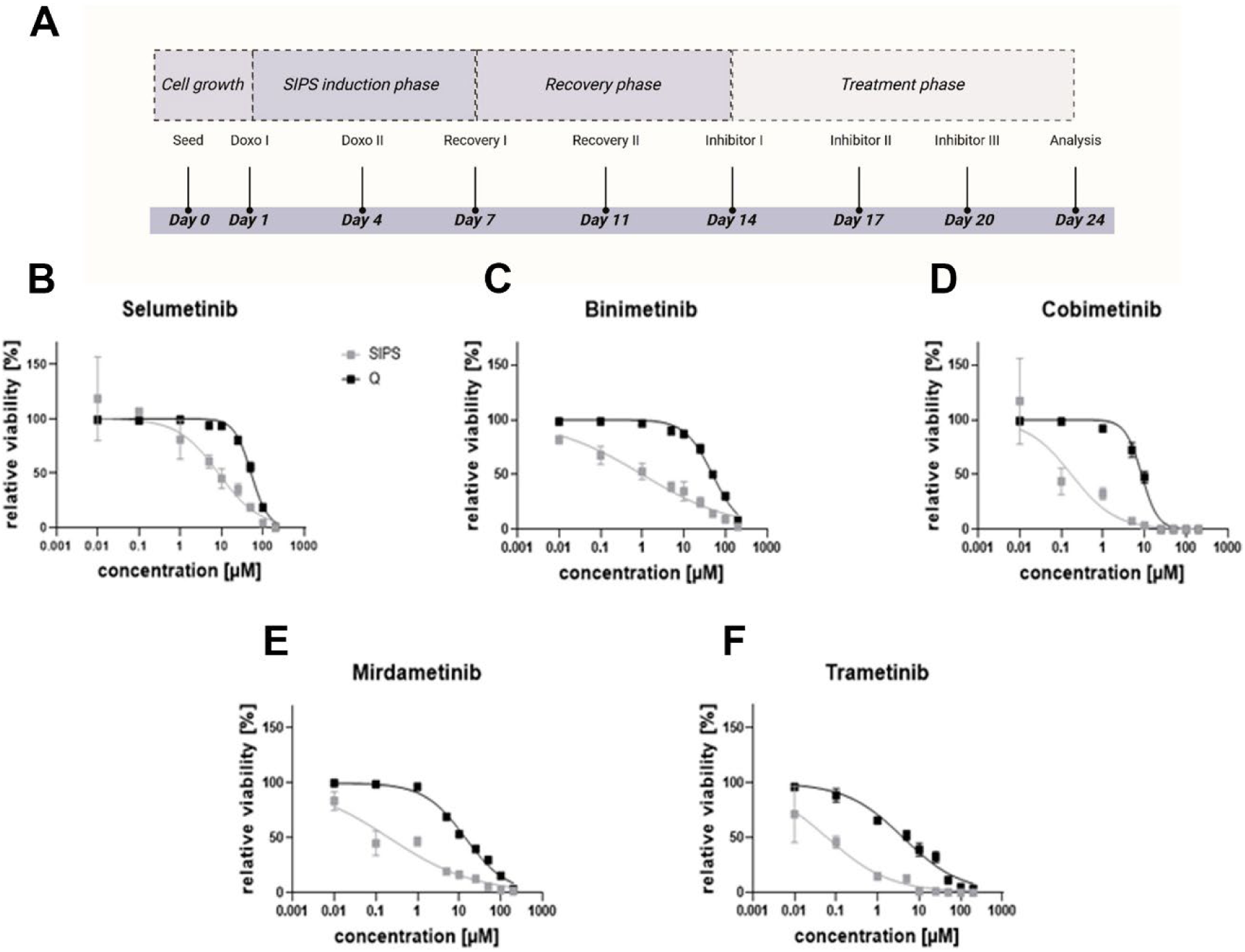
Senolytic effect of MEK inhibitors. (A) Schematic of evaluating the senolytic effect of MEK inhibitors in the stress-induced premature senescence (SIPS) model triggered by doxorubicin. (B-F) Viability plots of SIPS cells versus Quiescent Human Lung Fibroblasts in the presence of 5 MEK inhibitors: Selumetinib, Binimetinib, Cobimetinib, Mirdametinib, and Trametinib.

## Discussion

Here we report an innovative in silico approach to identify pathways and gene networks involved in making senescent human lung fibroblasts refractory to apoptosis and to identify potential drugs that can act as senolytic in this context. This analysis was based on re-using publicly available data and resulted in identification of a senescence signature that fits very well into current knowledge of the field, as outlined in detail below. In addition, it resulted in identification of MEK inhibitors as indeed senolytic in the context of in vitro lung fibroblast senescence.

### Senescence signature: cell cycle, and DNA repair and replication

Cell cycle arrest besides DNA replication and repair defects as represented in the KEGG pathways enrichment analysis of the obtained consensus senescence signature have been widely accepted as hallmarks of senescence [30, 31], also in vivo [32]. A large number of DNA repair genes are listed in downregulated signature, however, a study showed inhibiting two of the genes, BRCA1, and BLM (both are in the underexpressed signature) might contribute to the downregulation of most other DNA repair genes and eventually cause senescence [31]. Fanconi Anemia proteins are mostly involved in DNA repair processes, and their overexpression has been shown to counteract oncogene-induced senescence [33, 34]. Proliferating cell nuclear antigen (PCNA) is one of the downregulated genes that encodes a protein that interacts with a large network of proteins involved in DNA replication and repair and its downregulation probably impacts a large number of associated processes [35]. PCNA expression is inversely regulated by p21 which is congruent with our signature [36]. Similarly, nucleotide metabolism is changed due to decrease of RRM1 and RRM2 which provide the nucleotide pool for DNA replication. Surprisingly, RRM2B is overexpressed and probably provides basal levels of deoxyribonucleoside triphosphates (dNTPs) for the remainder of DNA repair activity in senescent cells similar to hypoxic cells, preventing their death [37]. Elevated expression of RRM2B in senescent cells [38] and a causal role of RRM2 depletion in senescence induction [39] were previously reported.

### Senescence signature: epigenetic modifications

GO overrepresentation analysis of the senescence signature as demonstrated in the Results section represent epigenetic changes and chromatin remodeling taking place in senescent lung fibroblasts. GO biological processes of “chromatin organization”, “positive regulation of histone H3-K4 methylation” and “negative regulation of histone H3-K9 methylation” are enriched in the downregulated signature. Both DNA methylation by downregulation of DNMT1 and DNMT3B and DNA demethylation by downregulation of TET1 and TET2 are at least transcriptionally decreased according to the consensus senescence signature. Consistently, it was shown that knockdown of DNMT1 triggers premature senescence, however, DNMT3B has been reported to be elevated at the protein level in senescent fibroblast cells of lung origin [40]. This controversy may arise from the post-transcriptional regulation and metabolism of DNMT3B which needs to be investigated. The authors further reported that congruent with their findings, replicative senescent human cells characteristically exhibit widespread DNA hypomethylation and focal hypermethylation [40]. Global DNA hypomethylation has been shown to be induced by the impaired function of BRCA1 through regulation of DNMT1 expression [41], therefore BRCA1 decrease as it is observed in the consensus signature might be regarded as one of the main culprits of large-scale chromatin remodeling and consequently senescence induction [42].

SETD2, SETMAR, and EZH2 are the genes that encode proteins with histone methyltransferase activity, and they are all in the underexpressed signature.

CBX1, CBX2, CBX5, ASF1B, SMCH1, High-Mobility Group genes (including HMGB1, HMGB2, HMGB3, and HMGN2) and condensin/cohesion complex genes (condensing I subunits including SMC2, SMC4, NCAPD2, NCAPH, and NCAPG; condensing II subunits including SMC2, SMC4, and NCAPG2; cohesion subunits including SMC1a and RAD21) are other genes that are all downregulated and are involved in epigenetic control or remodeling of the chromatin. CBX1 encodes Heterochromatin Protein 1-Beta (HP1 beta), and CBX5 encodes Heterochromatin Protein 1-Alpha (HP1 alpha). It is shown that high-level HP1 proteins, which are involved in heterochromatin formation, are not required for evolving the senescence-associated heterochromatin foci (SAHF) and other hallmarks of senescence, such as the expression of senescence-associated β-galactosidase activity. However, how its decrease might contribute to SAHF needs to be examined [43]. Similarly, ASF1B which is a histone chaperone has been argued to have failed to induce SAHF stems, at least in part, from its failure to bind HIRA [44], still, whether endogenous downregulation of this gene in senescent lung fibroblasts has anything to do with SAHF needs to be investigated.

SETMAR enhances nonhomologous end-joining (NHEJ) DNA repair by di-methylating H3K36 at DNA double-strand breaks (DSBs) [45], hence its downregulation might impair DSB repair in senescence. Similarly, SETD2 is reported to be responsible for methylation of H3K36, however, it only confers its tri-methylation, not mono-or di-methylation [46, 47]. To our knowledge depletion of SETMAR and SETD2 genes in senescence have not been investigated properly yet, this necessitates further studies to decipher their effect on chromatin organization and the gene expression regulation or DNA damage or replication responses in senescent cells.

EZH2 is the catalytic core protein of polycomb repressor complex 2 (PRC2) which transfers methyl to histone H3 at the lysine 27 (H3K27). Furthermore, CBX2 and SCMH1 which are among the core proteins in polycomb repressor complex 1 (PRC1) participate in transcriptional repression of the downstream genes by targeting the tri-methylated H3K27 residue (H3K27me3) after the methylation by PRC2 [48, 49]. It is shown that the downregulation of EZH2 promotes senescence through the initiation of a DNA damage response independent of the loss of H3K27me3 and then by the overexpression of the p16 (CDKN2A) which is induced by the subsequent depletion of H3K27me3 [50]. In another study, repression of H3K27 methylation with the EZH2 inhibition provided survival benefit and conferred chemotherapy resistance in testicular germ cell tumors while induction of H3K27 methylation with the histone lysine demethylase inhibition resulted in increased cisplatin sensitivity [51]. Interestingly, EZH2 downregulation has also been reported as one of the main drivers of UVB-induced senescence program in dermal fibroblasts [52]. Similarly, the SCMH1 transcription defect has been shown to induce premature senescence in mouse embryonic fibroblasts [53]. In addition, depletion of PHF19 which regulates PRC2 function has been shown to strongly reduce the global levels of H3K27me3 [54]. In a distinct study, it is demonstrated that PHF19 depletion causes an increase in the accumulation of H3K27me3 in genes associated with differentiation (not all genes) of hematopoietic stem cells [55], hence it can be deduced that PHF19 depletion is a cellular mean for abortion of differentiation capability of cells, probably the same thing as what happens in senescence.

Consistent with what gene ontology analysis of the signature denotes on the enrichment of the downregulated genes in the biological process of negative regulation of histone H3K9 methylation, it is previously reported that the depletion of PHF2, which is a histone demethylase, leads to the accumulation of H3K9me3 and causes an increase in heterochromatin-related marks at global levels [56]. Interestingly, TET1-depleted lung cancer cells demonstrated a reduction of di-methylated histone H3K9 (H3K9me2). This depletion sensitized the cells for therapy-induced senescence. The effect was shown to be independent of the defect in the demethylation of CpG islands [57]. Regarding the aforementioned studies on the role of chromatin remodeling players we hypothesize that in senescence, histone H3K9s proceed toward more tri-methylation and less di-methylation. Previous studies support this hypothesis. One study manifested that H3K9me3 mediates epigenetic regulation of senescence [58]. Another study approved the global decline of H3K9me2 in senescent cells. This decline contributed to the escalation of inflammatory responses which resembled characteristics of SASP [59]. It has been suggested that reduced H3K9me2 enhances the binding of NFκB and AP-1 (activator protein-1) transcription factors at specific inflammation-responsive genes to augment proinflammatory stimuli [60]. Therefore, epigenetic changes can elucidate the underlying event which gives rise to the prominent characteristics of senescence including SAHF and SASP. It has to be taken into account that histone methylation profiles may differ on a local scale compared to a global scale. As it is shown that the loss of Lamin B Receptor (LBR) in fibroblasts of lung origin, reduced H3K9me3 heterochromatin marks in lamina-associated domains (LADs) region of DNA and heightened transcription of LAD-associated repetitive elements. The loss led to senescence in the cells [61].

Among downregulated non-histone proteins that modify the chromatin organization, two proteins of HMGB1 and HMGB2 have been studied in terms of their association with senescence. HMGB1 and HMGB2 bind to distinct genomic loci, however, target loci of both HMGB1 and HMGB2 are usually upregulated once relieved of HMGB1 and HMGB2 binding, respectively. The loss of HMGB1 has been demonstrated to trigger p21 activation or SASP induction, in contrast, the loss of HMGB2 does not induce senescence but triggers senescence-induced CTCF clustering and loop reshuffling which act as a primer for the ensuing senescent program [62, 63]. The only gene that has been proven to affect chromatin organization and was upregulated in the consensus signature is TERF2IP. TERF2IP overexpression in senescence has been shown to protect critically short telomeres and its depletion in senescent cells that return to growth upon p21 inhibition caused higher levels of apoptosis [64]. TERF2IP interacts directly with H3/H4 histone tetramers and mediates chromatin opening required for new gene expressions in senescence [65]. In addition, TERF2IP has been demonstrated to repress histone gene expression and contribute to the global downregulation of histones in senescent cells [66]. In a different study, the depletion of total histone content in senescent cells was shown to be dependent on lysosomal-mediated proteolytic processing [67]. The nuclear leakage of chromatin is proposed to be caused by Lamin B1 (LMNB1) nuclear membrane protein loss in senescence and thereafter ubiquitinated chromatin is carried by adaptor protein, p62, to lysosomes for digestion [67]. Interestingly the consensus signature direction of gene expression is in agreement with these phenomena including LMNB1 and LMNB2 downregulation and p62 and cathepsins (CTSB and CTSS) upregulation.

### Senescence signature: Acidity, proteostasis and calcium regulation

Cytosolic decrease in acidity has been proposed as one of the drivers of apoptosis [68]. A study demonstrated that cardiac glycosides probably by induction of a dramatic drop in cytosolic pH of senescent cells contribute to apoptosis, so-called senolysis, of these cells [69]. In a recent study glutaminolysis by senescent cells has been presented as these cells’ defense tool against apoptosis which increases intracellular pH or at least counteracts pH drop by producing ammonia [70]. GLS gene expression is not increased in our consensus signature, however, analysis of gene ontology (Fig. S1) shows a reduction in intracellular organelles’ pH by an increase in genes involved in proton transport mainly by V-ATPases (ATP6AP1, ATP6AP2, ATP6V0A1, ATP6V0B, ATP6V0C, ATP6V1C1, ATP6V1E1, ATP6V1F, ATP6V1G1, ATP6V1H), and chloride transport by Chloride Voltage-Gated Channel 3 (CLCN3) [71], this theoretically accompanies the increase in cytosolic pH. It was shown that the granulocyte colony-stimulating factor (G-CSF) by upregulating the V-ATPase delays apoptosis in neutrophils through abolishing cytosolic acidification during the early stages of apoptosis [72]. Also, the NHE1 channel (SLC9A1) which has been shown to elicit an anti-apoptotic effect by cellular outport of protons has been upregulated in the signature [73, 74]. Contrary to previous reports of lysosomal alkalinization in senescent cells which had been deduced from observations in aging-related studies, upregulation of V-ATPases and CLCN3 beside MCOLN1 (TRPML1) [75] and OSTM1 might contribute to increased acidity of lysosomal lumina at least from transcriptional perspective. It must be noted that the increase in expression of genes involved in the regulation of intracellular pH might just be a compensatory response to the disrupted permeability of the lysosomal membrane that contributes to lysosomal leakage of protons in senescent cells. This cross-compartmental maintenance of pH balance rescues the senescent cells from apoptosis [76]. Moreover, GO biological processes involved in phagosome acidification and maturation have been enriched in the upregulated signature. Further digging into the signature, we found out that the expression of SQSTM1 (P62), MAP1LC3B (LC3B), GABARAPL2, and ATG4A which are the genes involved in autophagosome formation and biogenesis are elevated. Intriguingly proteasome inhibition has been demonstrated to induce a similar phenotype and rapidly elevate the expression of p62, LC3B, and ATG4A regardless of activation of other compartments of autophagy [77]. This has been shown to have a survival benefit for the cells even before the commencement of autophagy [77]. In this sense, distinct proteasome subunits were downregulated in differential gene expression datasets of replicative and RAS-induced senescence which we employed in this study; however, none were included in the consensus signature. In replicative senescence proteasome subunits of PSMA4, PSMA7, PSMB1, PSMB7, PSMC3, PSMD11, PSMD14, PSME3, PSMG1, and PSMG3 were downregulated, and in RAS-induced senescence, PSME1 and PSMB9 were downregulated. Some of the subunits of proteasome were overexpressed in RAS-induced senescence but none in replicative senescence (data not shown). The reduced proteasomal activity was previously reported in replicative senescence [78], however, whether the response to proteotoxic stress is similar in differently triggered senescent cells, has to be investigated. In addition, there are some conflicting results over the autophagy activation in senescent cells. Some of the most comprehensive studies confirm the activation of autophagy in fully executed RAS-induced senescence [79, 80]. Surprisingly, besides the autophagy-related genes that we noted, ATG101, ULK3, and ATG4D were exclusively overexpressed in datasets of RAS-induced senescence that we employed, but not in other forms of senescence. Studies that are denying the activation of autophagy in senescence mostly studied the role of autophagy in senescence induction rather than evaluating the effect on fully senescent cells. We speculate that impaired autophagy, as it happens during aging or direct stress to the compartments of autophagy, is a risk factor for senescence induction [81–83]. However, it seems when a cell is fully senescent, activation of different forms of autophagy including chaperone-mediated and selective autophagy might be crucial for confronting proteotoxic stresses and maintaining cell survival [84]. TPCN1 which is encoding a two-pore channel and TRPML1 regulate intracellular Ca^2+^ through endolysosomal system [85, 86]. Overexpression of these two genes probably increase calcium release to cytosol. In line with the increased cytosolic calcium and consequent altered signaling, TRPML1 expression has been shown to upregulate autophagic genes, increase autophagosomal biogenesis, and autophagosome-lysosome fusion [85]. Beside TPCN1 and TRPML1 two other genes of ITPR1 and TRPA1 are also included in the consensus upregulated signature which is enriched for GO molecular function of “calcium-release channel activity” (Fig. S1 and Dataset S1. Sheet3). ITPR1 is responsible for the release of calcium from the endoplasmic reticulum upon activation by cytosolic Ca^2+^ and IP3. TNF-α signaling and subsequent increase in PKC-α (PRKCA), have been demonstrated to contribute to ITPR1 (IP3R1) overexpression and an increase in cytosolic Ca^2+^ release. In the study, RNAi targeting PKC-α inhibited TNF-α-induced IP3R1 overexpression and restored cytosolic Ca^2+^ concentration [87]. The increased calcium release activates a number of pathways which consequently leads to the development of some of the phenotypes of senescence, like the inflammatory ones [88]. IP3R1 knockdown has been shown to promote escape from OIS [89]. Cytosolic calcium release has also been discussed in many studies as the driver of apoptotic cell death [90]. However, simultaneous activation of unfolded protein response (UPR) mediated by ERN1 (IRE1) has been shown to protect cells from apoptosis as ERN1 knockdown led to apoptosis by IP3R1-mediated calcium release [91]. UPR is the cell reaction to the accumulation of misfolded proteins above a critical threshold, in order to restore the defect in protein homeostasis [92]. GO analysis confirms enrichment of the upregulated signature in the biological process of “IRE1-mediated unfolded protein response”. IRE1 overexpression was previously reported to be implicated in promoting HRas-induced senescence [93]. We hypothesize that in ER of senescent cells, UPR and calcium homeostasis are finely balanced so that the outcome is evasion from apoptotic cell death despite the proteostasis fragility in senescent cells [94]. Transcriptional increase of the co-chaperone, DNAJB9, the downstream of ERN1-mediated UPR activation [95] not only regulates ERN1 activation in a feedback manner [96] but also has been shown to exert an antiapoptotic effect by physical interaction with p53; therefore, it is one of the possible routes how ERN1 activation might abolish apoptosis [97]. We speculate that ERN1 upregulation in the signature is the consequence of demethylation of histone H3K27-me3 at the promoter region of ERN1 enforced by EZH2 downregulation [98] in senescent cells. ER calcium depletion by genetic ablation of sarco/endoplasmic reticulum Ca^2+^-ATPase (SERCA2) or by its inhibition using thapsigargin has been demonstrated to induce senescence phenotype [99]. Intriguingly, thapsigargin not only simulates ER stress and UPR, but also enhance autophagosome formation, similar to our findings through the consensus signature of senescence. However, thapsigargin probably due to the vast alteration of intracellular calcium signaling disrupts autophagosome-lysosomal fusion and ultimate autophagy flux [100]. A more thorough mechanistic evaluation of the senescence-mimetic effect of thapsigargin may help enlighten the root causes of transcriptional changes in senescence.

### Senescence signature: Lipid metabolism

The other concurrently activated pathway that took our attention while interpreting the consensus signature is the increased metabolism of sphingolipids which is enriched in both KEGG pathways and GO biological processes. KDSR upregulation is at the forefront of the pathway involved in the de novo production of ceramide and sphingosine [101]. The balance between the accumulation of ceramides by overexpression of GBA and CERS2, and the accumulation of sphingosine and sphingosine-1-phosphate (S1P) by upregulation of ACER2 presumably have led the cells to preferably go toward senescence state rather than apoptosis. It is shown in a study, that ACER2 upregulation increases the levels of both sphingosine and S1P while decreases the levels of ceramides in cells. They have manifested that a moderate upregulation of ACER2 in an interplay with p53 signaling gives rise to cellular senescence by elevating S1P, a pro-proliferative and pro-survival bioactive lipid [102]. Consistently, knockout of NSMAF (FAN), which is downregulated in our signature and was reported to mediate tumor necrosis factor-induced activation of neutral sphingomyelinase, has been demonstrated to reduce cellular ceramide level by inhibiting CD40 ligand-induced sphingomyelinase stimulation. The changes eventually abrogated apoptosis of human fibroblasts [103]. In this regard, cleavage of ceramide into sphingosine by acid ceramidase (ASAH1) has been reported to potentiate resistance of senescent cells against senolytics. ASAH1 was not included in our consensus signature, however, surprisingly, we noticed that it has been upregulated in all datasets of RS and IRIS (not OIS) [104]. The other overexpressed gene involved in the metabolism of sphingolipids is prosaposin (PSAP) which facilitates the catabolism of glycosphingolipids. Overexpression of this gene was demonstrated to mediate inflammatory responses [105] which may associate with SASP. Furthermore, ADIPOR1 overexpression which is linked with ceramide catabolism might have contributed to skewing the balance toward an enhanced formation of S1P, ultimately exerting an antiapoptotic effect [106].

### Senescence signature: ERK

GO analysis of the molecular functions of the overexpressed signature also indicates enrichment of “MAP kinase phosphatase activity” via 3 Dual-specificity phosphatases (DUSP3, DUSP4, and DUSP6) which dephosphorylate active mitogen-activated protein kinases (MAPKs) including ERK1/2. It is shown that DUSP4 ablation of gene expression or expression of the phosphatase-resistant ERK2 postpones replicative senescence [107]. Another study reported reduced phosphorylated ERK representing the impaired ERK activation in senescence and pointed to the increased DUSP6 expression as the culprit of this defect. Their investigation has also shown that the loss of DUSP6 activity in senescent cells restores ERK1/2 phosphorylation and promoted cell proliferation [108], however, they have not studied any other hallmarks of senescence. We hypothesize that the DUSP6 transcriptional activation is probably the apoptotic-rescue response of the prolonged overactivation of ERK1/2 which if persisted could lead to cell death [109, 110].

### Senescence signature: Mitochondrial homeostasis

The consensus signature also reflects changes in mitochondrial homeostasis. It has been indicated that while ATP production through glycolysis increases in senescent cells, the overall production of ATP mainly through impaired oxidative phosphorylation is reduced [111–113]. However, it turns out that in the signature, mitochondrial genes involved in oxidative phosphorylation, which is overrepresented in KEGG pathways, are upregulated. The upregulated genes including ND1, ND2, ND3, ND4, ND4L, ND5, COQ10B, and CYTB are enrolling in the respiratory electron transport chain. This overexpression, besides the underlying increase in mitochondrial gene copy number in senescent cells (due to increased mitochondrial biogenesis), might be representative of the effort of the cells to repair the defect in ATP production through oxidative phosphorylation. By exploring the signature, we found out that the directions of regulation of some of the genes are in agreement with the notion of the failure in the electron respiratory chain. One of the genes is MTFR2 (DUFD1) which is involved in mitochondrial fission, and its expression is ameliorated in the consensus senescence signature. It is shown that its Inhibition severely impairs O2 consumption which is indicative of its importance in mitochondrial respiration [114]. Similar to cytosolic and ER calcium homeostasis that we discussed mitochondrial calcium homeostasis has a crucial role in the induction of cellular senescence. Loss of FUS1, which is a DNA/RNA-binding protein with multiple functions, in a study (also downregulated in the signature) caused inefficient accumulation of Ca^2+^ in mitochondria and decreased respiratory reserve capacity [115]. MCUB encodes a protein that negatively regulates the activity of the mitochondrial inner membrane calcium uniporter (MCU) [116]. MCU mediates calcium uptake into the mitochondria [117]. Therefore, the downregulation of MCUB in the senescence signature must have brought about the increase in mitochondrial calcium level [118]. It has been shown that during OIS, ITPR2 (most probably the same role for IP3R1) triggers calcium release from the endoplasmic reticulum, followed by mitochondrial calcium accumulation through MCU channels. Mitochondrial calcium accumulation has been claimed to lead to a subsequent decrease in mitochondrial membrane potential, reactive oxygen species accumulation, and senescence perse [89].

SLC25A40 (Solute Carrier Family 25 Member 40) which is a paralog of the gene SLC25A39, is probably involved in the mitochondrial import of glutathione. Mitochondrial glutathione by detoxifying ROS is required for the activity and stability of proteins that contain iron-sulfur clusters, involved in respiratory electron transport [119]. Thus, downregulation of this gene in the signature put mitochondrial respiration more prone to ROS-induced damage. CISD1 (MitoNEET) is another gene whose expression is enhanced according to the senescence signature. MitoNEET is a protein that contains a 2Fe–2S cluster and is located at the outer mitochondrial membrane. It is involved in the regulation of electron transport and oxidative phosphorylation [120]. Also, it regulates cellular iron and ROS homeostasis [121]. The expression brings about resistance to iron-mediated cell death (ferroptosis) [122] which is congruent with the previous reports of increased resistance to ferroptosis in senescent cells [123]. Looking through the signature, sideroflexin 2 (SFXN2) is another downregulated gene that is associated with mitochondrial transmembrane transport. knockout (KO) of SFXN2 in a study has been shown to increase mitochondrial iron content, which is suggested to be associated with decreases in the heme content. The authors claimed the abnormal iron metabolism impaired mitochondrial respiration in SFXN2-KO cells which can be similar to the phenomena in the senescent cells [124]. We hypothesize that the increase in expression of the compartments of the mitochondrial electron transport system, concurrent with the flawed oxidative phosphorylation might have given rise to the increase in reactive oxygen species (ROS) generation as it was previously shown that inhibiting the expression of the Rieske iron-sulfur protein of complex III (containing 2Fe–2S cluster) or its pharmacological inhibition elevated ROS level and decreased ATP and consequently was sufficient to trigger senescence [125]. To deal with this high-level ROS production without bringing the cell into apoptosis, cells seem to utilize thiol-specific peroxidases to protect the cell against oxidative stress by detoxifying peroxides [126]. Peroxiredoxin 2 (PRDX2) is among the family of these genes which is upregulated in the signature and undertakes this role [127]. It is probably one of the mechanisms in senescent cells under our study which provokes resistance against apoptosis due to the high level of ROS.

### Senescence signature: RNA splicing

15 genes related to RNA splicing have been represented in the underexpressed signature; None have been upregulated. GO biological process of “regulation of RNA splicing” has been overrepresented (Dataset S1. Sheet3). Genetic depletion of SRSF2 [128, 129], SRSF3 [130, 131], HNRNPA1 [132], HNRNPD [129], HNRNPU [133] and TRA2B [134] have previously been shown to bring about or associate with senescence through regulating different pathways including p21 and/or p53 [128, 130, 131, 133, 134].

### Key elements of senescence signature conferring pro-survival effect

The senescence inductive effect of p21 overexpression which is a cell cycle inhibitor has been evident for quite a long time, however, its role in guaranteeing the viability of senescent cells has also been manifested recently which is consistent with our findings analyzing CMap data [135]. We previously noted the hypothetical role of PRDX2 in protecting senescent cells against high-level mitochondrial ROS [127]. Our transcriptional connectivity findings also support this pro-survival role.

The computational analysis of RNAi LOF perturbation in cancer cell lines that mimic senescence mechanisms of survival, revealed JUNB, SQSTM1, LIF, and AREG as critical genes in senescent cell survival. JUNB expression has been shown to be responsible for the induction of a large portion of the senescence phenotype including cell cycle arrest [136] and proinflammatory traits [137]. JUNB overexpression has been indicated to be protective against ER stress (which is observed in senescence) through activating AKT pathway [138]. It was claimed by a study that JUNB has repressive effect on expression of p62 [139], however, at least in senescence this repressive effect has been overcome by other factors causing overexpression of this gene. P62 has been demonstrated to be prosurvival factor in senescence through the NF-κB pathway, and its knockdown, incongruent to our speculation, has increased apoptotic cell death of senescent cells [140]. Moreover, the deubiquitinating enzyme USP20 which is also overexpressed in the consensus senescence signature, by stabilizing SQSTM1 contributes to TNFα /NF-κB-mediated cell survival [141]. A positive feedback loop between p62 expression and NF-κB activation has been reported elsewhere [142].

Amphiregulin (AREG) is overexpressed as a consequence of mitochondrial dysfunction [143] and its expression has been shown to be a part of a pathway that leads to senescence in BRAFV600E transduced melanocytes; The inhibition of AREG caused a reduction in promyelocytic leukemia protein (PML), p21, and gamma-H2AX protein expression, characteristics of senescence, suggesting an association of the senescence induction process to amphiregulin [144]. However, whether its decrease in senescent cells promotes cell apoptosis has not been investigated and remains to be determined. Our current knowledge of leukemia inhibitory factor (LIF) in regard to senescence is still insufficient. It contributes to radio-resistance in nasopharyngeal carcinoma [145] which could indicate an anti-apoptotic role, while its induction by TGFβ is required for TGFβ-mediated overexpression of p21 and consequently cell cycle arrest [146] as a hint towards a role in senescence induction or maintenance.

All these reports are well in line with our bioinformatic findings using transcriptional connectivity and genetic perturbation sensitivity as well as the gene set enrichment and literature-based interpretations of the whole consensus senescence signature.

Moreover, we bioinformatically identified selumetinib, which is a MEK inhibitor, alongside other MEK inhibitors as a class of senolytics. MEK inhibitors have been claimed to induce autophagy flux and diminish the abundance of p62 in BRAF and RAS mutant cancer cell lines [147]. A previous study has shown that MEK inhibitors impair autophagosome-lysosome fusion and generally autophagy flux when simultaneously used with senescence inducer in RAS mutant rat embryonic fibroblasts thereby restraining cell survival [148].This must be noted that senescence program was not fully implemented at the time of the administration of the MEK inhibitor. Surprisingly, such activity was indeed observed in adult human lung fibroblasts in which senescence was definitive as predicted by our computational model. MEK has been shown to activate Heat Shock Transcription Factor 1 (HSF1) to enhance the proteostasis in cells by increasing the chaperoning capacity required for selective autophagy and confronting the proteotoxic stress [149, 150]. Surprisingly, HSF1 inhibition has been shown to impair aggregate-induced autophagosome formation and p62-mediated proteostasis [151]. Therefore, we speculate that MEK inhibitors might exert their senolytic effect through alterations in p62 and molecular chaperones including heat shock proteins which are at the downstream of MEK/HSF1 axis. Accordingly, knockdown of p62 [140] and inhibition of HSP90 [152], a molecular chaperone, have been manifested to induce apoptosis in senescent cells. Further, we propose p62 and molecular chaperones as potential target hubs for future development of potent senolytics. Moreover, the MEK/ERK axis has been implicated in promotion of ERN1-mediated OIS [93]. As we noted above, ERN1-mediated UPR and calcium signaling, together, participate in provoking resistance against apoptosis. However, to our knowledge whether MEK/ERK axis inhibition can induce apoptosis through this pathway has not been explored nor in general nor in senescence.

There are some limitations to our study. As our study was a vertical propagation of steps each being performed one after another, we were losing data in each step depending on the availability of the shared data in the databases. A number of the data losses include but are limited to: not utilization of a few queried genes by CMap in the calculation of NCSs; CCLE gene expression profiles did not contain all genes in the consensus senescence signature which might have affected our cell line correlation analyses; Cell lines were not identical between CCLE and sensitivity datasets, thus we used merely the shared ones for the analyses. Sensitivity data related to the majority of the small-molecules and gene LOF perturbations were not available, also sensitivity data of gene overexpression interventions were fully lacking. Validation of CMap-based prediction of senolytic effect due to the rareness of well-identified senolytics which exert their impact on senescent lung fibroblasts, was not possible for our team. Moreover, subjective curation of the anti-apoptotic signature alongside the evidence-based ranking and extracting of the genes might have caused us to falsely classify or lose some important genes which are not well-studied up to the time of this study. Still, since our final gene signatures were quite large, we are optimistic that results are not overly skewed.

One of the things that might have flawed our interpretations through the CMap data, which was also warned by the CMap team, is that gene overexpression does not always enhance the function of the targeted gene and pathways, sometimes due to the cellular compensatory responses. The last concern we would like to mention is the Groups’ sample size variation in the permutation tests which might have affected the results.

### Conclusion

To sum up, in the current study we identified a consensus senescence signature specifically related to fibroblasts of lung origin and semi-manually extracted its genetic module which functions as the anti-apoptotic compartment of the senescence phenotype. Afterward, with this assumption that we might be able to employ datasets related to cancer cell lines which are quite large nowadays, and extrapolate them to other fields, we tried to take advantage of these valuable deposited data to discover new senolytics which are now hotspots of cancer and aging sciences. In this way, using CMap transcriptional connectivity data we filtered several compounds and genetic interventions that could at least transcriptionally counteract the apoptosis-resistance signature which is exploited by the senescent cells to survive. Ultimately, by transcriptional connectivity-based analyses alongside the evaluation of cell survival response (all in cancer cell lines) in exposure to the compounds or genetic interventions, we suggested a MEK inhibitor and p62 or JUNB depletion as potential inducers of apoptosis in senescent lung fibroblasts. The senolytic effect of a range of MEK inhibitors was confirmed in vitro.

## Methods

### Obtaining the consensus senescence signature in fibroblasts of lung origin

In the beginning, we gathered 19 RNA-sequencing gene expression datasets (from 10 different studies) of senescence induced by different triggers in IMR90 and WI38 cells (both fibroblasts with lung origin) from GEO. Three datasets of oncogenic RAF, etoposide and doxorubicin induced senescence in lung fibroblasts were removed because we could find just one dataset for each, making us unable to validate its accuracy through comparison with their counterparts. Remaining datasets were related to Ras, replicative exhaustion, and Irradiation induced senescence. Differential gene expression analysis was performed by the Grein app. All datasets are analyzed through the same method by the app (TMM normalization). Using principal component analysis (PCA), 2 remote datasets were removed, leaving 14 datasets. In addition, using hierarchical clustering, remote samples for differential expression analysis were removed (detailed metadata of the final used datasets and their samples are available in Dataset S1. Sheet1). To reach a unified signature from these 14 datasets, which does not exclude any gene with a critical role in the core senescence phenotype, instead of conducting a conventional meta-analysis, we innovated our own approach. Here is the method (python script is available in the supplementary files):

For including any gene in the core senescence signature, the gene should have been expressed in the same direction in all datasets or not significantly (FDR adjusted p-value cut-off strictly set at 0.1) inversely expressed in any dataset. Moreover, the gene should have been significantly differentially expressed (FDR adjusted P-value cut-off= 0.05) in more than 75% of the datasets.

### Deriving the anti-apoptotic signature from the obtained senescence signature

Genecards’ term-scoring system and Gene Ontology (GO) annotations alongside literature reviewing were the tools we utilized to find out the core gene system resisting apoptosis in senescent cells. The “apoptosis” term was searched in the Genecards database (searched at 2022/01/22) [153]. The output was a list of genes that were ranked based on their association with the “apoptosis” term which was obtained by Genecards’ algorithmic searching through scientific literature and bioinformatics databases. Genes with a score of more than 1 were filtered (we excluded the rest of the genes because the manual curation of the whole genes had minimal benefit regarding its time cost). Moreover, genes annotated for GO of “Regulation of Cell Death”, which were obtained from the AmiGO web application, were appended to the previously filtered genes and were included in our manual evaluation and curation of the genes. Around 300 genes related to apoptosis and cell death were extracted. Afterward, we manually searched the publications for the relation of these genes with apoptosis regarding the direction of the expression (more than 1500 articles were read). To find out the association of these genes with apoptosis we searched if their modulation in the same direction as expressed in the senescence signature has shown an anti-apoptotic effect or raised resistance against chemotherapeutics or if their inverse modulation could lead to an apoptotic effect in different studies (most studies were related to cancer cell lines). Overall, 116 genes (57 downregulated and 59 upregulated) were extracted. The table of the anti-apoptotic signature (also we call it the signature of interest throughout the study) and the references based on which we established our decision are available in Supplementary file 1.

### Overrepresentation analysis of the signatures

Gene signatures enrichment analyses for KEGG pathways, GO biological processes, and GO molecular functions were performed in R (version 4.2.0) using ClusterProfiler library (v4.4.1) [154]. Significance was reported by p-value, adjusted by Benjamini-Hochberg FDR correction. Enrichment analysis for MSigDB hallmark gene sets (2020) was conducted using EnrichR library [155].We used p-value for the analysis of significance.

### Cell lines with the most similar and dissimilar gene expression profiles regarding the senescence signature and the anti-apoptotic signature

The dataset of the standardized gene expression profiles of Cancer Cell Line Encyclopedia (CCLE) [156] was downloaded from Harmonizome [157] to find out the most positively and inversely (most negatively) correlated cell lines with the senescence signature and the anti-apoptotic signature of interest. For analysis of the correlation, we used the point biserial correlation function from the SciPy statistical tool (version 1.8.0) in python (3.9.12). Upregulated genes were annotated as “1” and downregulated ones were annotated as “-1”. Cut-offs for correlation scores were determined differently and separately explained in the dedicated sections. The p-value for significance was set at 0.05.

### Connectivity map of the anti-apoptotic signature

The anti-apoptotic signature was submitted into the CLUE Query platform to obtain the Connectivity Map (CMap) of the small-molecules and genetic perturbations [10]. Genes and compounds were ranked using the mean of the normalized connectivity scores (NCSs) of the genes or compounds in their different perturbations (different doses, perturbation time, perturbed cell lines, and perturbation-type for the genes). We excluded perturbations in cell types whose gene expression signatures were negatively correlated with the anti-apoptotic signature (point-biserial correlation coefficient <-0.2 and p-value<0.05) from the calculation of the mean of the NCSs (MCS). Compound entities with less than 10 perturbations and a median TAS score of less than 0.1 were excluded from further rankings and selections. We used CMap to identify and rank the contribution of each gene in the anti-apoptotic signature to the development of the whole anti-apoptotic module. For analysis of the transcriptional effect of the inhibition of the upregulated genes, we measured the MCSs of the CRISPR loss of functions (which was annotated as “trt_xpr” in the perturbation-type column of the dataset) and the MCSs of the short hairpin RNA (shRNA) loss of functions (we only used NCSs of the consensus signature from shRNAs targeting the same gene which was annotated as “trt_sh.cgs” in the perturbation-type column of the dataset). For analyzing the transcriptional effect of the inverse modulation of the downregulated genes, we calculated the MCS of the gene overexpression (which was annotated as “trt_oe” in the perturbation-type column of the dataset). Gene entities of the same perturbation type with less than 4 perturbations and a median TAS score of less than 0.1 were excluded from the rest of the study analyses to reduce errors due to the low sample number and lack of reproducibility. Furthermore, the significance of the differences between the NCSs of the gene LOF (scores of perturbations with “trt_sh.cgs” and “trt_xpr” perturbation type, were pooled) and the NCSs of the gene GOF, was analyzed using the permutation test. The alternative hypothesis of the permutation test was set in a way that the mean of the NCSs of a set of gene perturbations corresponding to a single gene intervention that opposes the expression direction of that gene in our anti-apoptotic signature should have been less than the mean of NCSs of a set of gene perturbations corresponding to a gene intervention that its direction of regulation is similar with the one exists in the anti-apoptotic signature. P-values below 0.05 were considered to be significant.

### Analysis of drug and gene loss of function sensitivity in cell lines of similar gene expression profiles vs dissimilar ones

We evaluated the cell viability impact of the compounds and the gene loss of functions in cell lines with similar gene expression profiles (correlated cell lines) versus cell lines with the most dissimilar gene expression profiles (inversely correlated cell lines). The significance was measured using the permutation test. For compound perturbation, “drug sensitivity AUC (CTD^2)” dataset and for gene loss of function, “CRISPR (DepMap 22Q2 Public+Score, Chronos)” and “RNAi (Achilles+DRIVE+Marcotte, DEMETER2)” datasets were downloaded from the DepMap portal. The correlations of the CCLE standardized gene expression profile of each cell with senescence signature and anti-apoptotic signature were assessed and the cell lines with a correlation coefficient of >0.4 with both senescence signature and the anti-apoptotic signature were considered as correlated cells (p-value<0.05) and cell lines with the correlation coefficient of <-0.4 for both senescence signature and the anti-apoptotic signature were considered as negatively correlated cells (p-value<0.05)(correlation analyses were performed on cell lines which were shared between CCLE datasets and each of the sensitivity datasets). To avoid bias toward any specific lineage of cancer cell lines, we used at most the two most correlated (or inversely correlated) cell lines in each lineage. As the sensitivity of different cell lines to the same drugs varies in scale due to the different transcriptional contexts, we standardized the sensitivity data of each cell line prior to the permutation test to be able to compare the sensitivity data on a similar scale. Perturbations were considered to be more lethal in correlated cell lines than inversely correlated ones if the subtraction of the mean of the standardized sensitivity scores in the inversely correlated cell lines from the mean of the standardized sensitivity scores in the correlated cell lines yielded a negative value and the p-value of the permutation test was less than 0.05. In addition, to assure that the significance of the different effects of the drug in correlated cell lines vs inversely correlated cell lines is due to more lethality in correlated cell lines, not the more proliferation enhancing effect on inversely correlated cell lines, we evaluated replicate level viability data of the final selected compounds (Drug sensitivity replicate-level dose (CTD^2) was downloaded from DepMap portal) and those with the overall mean viability of more than 0.9 in both correlated and inversely correlated (not just one of them), were excluded.

### Other statistical and visualization tests and transformations

PCA was performed on log fold changes data of differential gene expression analysis using sklearn library (version 1.0.2), decomposition tool, and plotted using Plotly (version 5.6.0) in python. Standardization (z-score) was performed using sklearn library preprocessing tool. The number of repeats in all permutation tests was set at 10000. The sample size cut-off for analysis of significance using the permutation test was set at a minimum of 4 for each group.

### Cell isolation and culture

Human parenchymal lung fibroblasts were isolated and cultured as described previously [158, 159].

## Author Contributions

Conceptualization: ANR, JG; Software and data curation: ANR; Methodology: ANR, LMM, KH, IL, SS; Investigation: ANR, SS; Formal analysis: ANR, SS, IL, JG; Resources: LMM, KH; Writing – Original draft: ANR, JG Writing: Review & Editing: all authors have edited, read and approved the submitted manuscript; Supervision: ANR, JG.

## Acknowledgments

We gratefully acknowledge the Austrian Science Fund (Österreichischer Wissenschaftsfonds; FWF) grant DOI: 10.55776/P35268.

