## Supplementary file 1 for "Connectivity Map and perturbation-based sensitivity analysis identifies MEK inhibitors as senolytics in human lung fibroblasts"

| Upregulated genes in the senescence signature which their upregulation have been shown to have an anti-apoptotic effect or exhibit a chemoresistance effect or their downregulation might increase apoptosis in different studies | | | |
| --- | --- | --- | --- |
| Gene Symbol | Gene Full Name | References for | References against |
| CDKN1A | Cyclin Dependent Kinase Inhibitor 1A | (1-6) | |
| TNFRSF10C | TNF Receptor Superfamily Member 10c | (7-9) | |
| TNFRSF10D | TNF Receptor Superfamily Member 10d | (10) | |
| BCL2L2 | BCL2 Like 2 | (11) | |
| VEGFA | Vascular Endothelial Growth Factor A | (12) | |
| IL6 | Interleukin 6 | (13-18) | |
| SQSTM1 | Sequestosome 1 | (19-28) | |
| TIGAR | TP53 Induced Glycolysis Regulatory Phosphatase | (29, 30) | |
| ERN1 | Endoplasmic Reticulum To Nucleus Signaling 1 | (31-34) | |
| SHC1 | SHC Adaptor Protein 1 | (35-37) | (38-42) |
| PTRH2 | Peptidyl-TRNA Hydrolase 2 | (43, 44) | (45) |
| CSF2 | Colony Stimulating Factor 2 | (46-52) | |
| HK2 | Hexokinase 2 | (53-55) | |
| GDNF | Glial Cell Derived Neurotrophic Factor | (56-58) | |
| IGF2R | Insulin Like Growth Factor 2 Receptor | (59, 60) | |
| NRG1 | Neuregulin 1 | (61-64) | |
| CD44 | CD44 Molecule (Indian Blood Group) | (65, 66) | |
| MXD1 | MAX Dimerization Protein 1 | (67, 68) | |
| OSMR | Oncostatin M Receptor | (69, 70) | |
| AREG | Amphiregulin | (71-74) | |
| PRDX2 | Peroxiredoxin 2 | (75-79) | |
| BSG | Basigin (Ok Blood Group) | (80-82) | |
| CHMP5 | Charged Multivesicular Body Protein 5 | (83) | |
| SLC9A1 | Solute Carrier Family 9 Member A1 | (84-86) | |
| ANGPTL4 | Angiopoietin Like 4 | (87-89) | |
| ADIPOR1 | Adiponectin Receptor 1 | (90-94) | |
| JUP | Junction Plakoglobin | (95, 96) | |
| DUSP6 | Dual Specificity Phosphatase 6 | (97-99) | |
| PROCR | Protein C Receptor | (100, 101) | |
| MAP1LC3B | Microtubule Associated Protein 1 Light Chain 3 Beta | (102-105) | (106-108) |
| LIF | LIF Interleukin 6 Family Cytokine | (109-111) | (112) |
| AKR1B1 | Aldo-Keto Reductase Family 1 Member B | (113-115) | |
| GLRX | Glutaredoxin | (116-119) | |
| FANK1 | Fibronectin Type III And Ankyrin Repeat Domains 1 | (120, 121) | |
| RRM2B | Ribonucleotide Reductase Regulatory TP53 Inducible Subunit M2B | (122-125) | |
| DNAJB9 | DnaJ Heat Shock Protein Family (Hsp40) Member B9 | (126) | |
| IL12A | Interleukin 12A | (127, 128) | (129) |
| CXCL1 | C-X-C Motif Chemokine Ligand 1 | (130-132) | |

| | | | |
|---|---|---|---|
| JUNB | JunB Proto-Oncogene, AP-1 Transcription Factor Subunit | (133-138) | |
| CD9 | CD9 Molecule | (139-143) | (144) |
| DDRGK1 | DDRGK Domain Containing 1 | (145-148) | |
| ABCA1 | ATP Binding Cassette Subfamily A Member 1 | (149-151) | |
| GABARAPL2 | GABA Type A Receptor Associated Protein Like 2 | (152, 153) | |
| CTSS | Cathepsin S | (154-158) | |
| PSENEN | Presenilin Enhancer, Gamma-Secretase Subunit | (159, 160) | |
| CXADR | CXADR Ig-Like Cell Adhesion Molecule | (161, 162) | |
| DUSP4 | Dual Specificity Phosphatase 4 | (163-168) | |
| FNTA | Farnesyltransferase, CAAX Box, Alpha | (169-171) | |
| PLAT | Plasminogen Activator, Tissue Type | (172, 173) | (174, 175) |
| LUCAT1 | Lung Cancer Associated Transcript 1 | (176-181) | |
| CLN8 | CLN8 Transmembrane ER And ERGIC Protein | (182-185) | |
| CLCN3 | Chloride Voltage-Gated Channel 3 | (186-188) | (189, 190) |
| PSAP | Prosaposin | (191-196) | |
| CHST11 | Carbohydrate Sulfotransferase 11 | (197, 198) | (199, 200) |
| CPEB4 | Cytoplasmic Polyadenylation Element Binding Protein 4 | (201-205) | |
| LRPAP1 | LDL Receptor Related Protein Associated Protein 1 | (206, 207) | |
| STXBP1 | Syntaxin Binding Protein 1 | (208-210) | |
| TM7SF3 | Transmembrane 7 Superfamily Member 3 | (211, 212) | |
| TNFSF18 | TNF Superfamily Member 18 | (213-215) | (216) |

| Downregulated genes in the senescence signature which their downregulation have been shown to have an anti-apoptotic effect or exhibit a chemoresistance effect or their upregulation might increase apoptosis in different studies | | | |
|---|---|---|---|
| GeneSymbol | Gene_Full_Name | References for | References against |
| CASP2 | Caspase 2 | (217-221) | |
| CASP6 | Caspase 6 | (220, 222, 223) | |
| PARP1 | Poly(ADP-Ribose) Polymerase 1 | (224-228) | |
| CDK1 | Cyclin Dependent Kinase 1 | (229-235) | |
| TET2 | Tet Methylcytosine Dioxygenase 2 | (236-238) | (239-241) |
| CDKN1B | Cyclin Dependent Kinase Inhibitor 1B | (242-245) | |
| PCNA | Proliferating Cell Nuclear Antigen | (246, 247) | |
| BRCA1 | BRCA1 DNA Repair Associated | (248-251) | |
| PRKCA | Protein Kinase C Alpha | (252-255) | (256-260) |
| CCNB1 | Cyclin B1 | (235, 261-267) | (268) |
| CASP8AP2 | Caspase 8 Associated Protein 2 | (269, 270) | |
| EZH2 | Enhancer Of Zeste 2 Polycomb Repressive Complex 2 Subunit | (236) | (271-276) |
| NOD1 | Nucleotide Binding Oligomerization Domain Containing 1 | (277-280) | |
| SET | SET Nuclear Proto-Oncogene | (281, 282) | (283-288) |
| MLH1 | MutL Homolog 1 | (289-295) | |
| HNRNPA1 | Heterogeneous Nuclear Ribonucleoprotein A1 | (296-298) | (299) |
| ITGB3BP | Integrin Subunit Beta 3 Binding Protein | (300-303) | |
| SMAD4 | SMAD Family Member 4 | (304-309) | (309, 310) |
| MSH2 | MutS Homolog 2 | (311-316) | |
| JADE1 | Jade Family PHD Finger 1 | (317) | |
| TOP2A | DNA Topoisomerase II Alpha | (318, 319) | (320-322) |
| SRSF2 | Serine And Arginine Rich Splicing Factor 2 | (299, 323, 324) | |
| BARD1 | BRCA1 Associated RING Domain 1 | (325-327) | (328) |
| MCM2 | Minichromosome Maintenance Complex Component 2 | (329) | (330) |
| E2F2 | E2F Transcription Factor 2 | (331, 332) | |
| SETD2 | SET Domain Containing 2, Histone Lysine Methyltransferase | (333-335) | (336) |
| ID4 | Inhibitor Of DNA Binding 4, HLH Protein | (337-344) | |
| TJP2 | Tight Junction Protein 2 | (345-349) | |
| KLF2 | Kruppel Like Factor 2 | (350) | (351, 352) |
| SMARCA2 | SWI/SNF Related, Matrix Associated, Actin Dependent Regulator Of Chromatin, Subfamily A, Member 2 | (353-355) | (356, 357) |
| MCM3 | Minichromosome Maintenance Complex Component 3 | (358) | (359, 360) |
| FUS | FUS RNA Binding Protein | (361-363) | |
| SENP1 | SUMO Specific Peptidase 1 | (97, 364) | |
| HNRNPD | Heterogeneous Nuclear Ribonucleoprotein D | (365-367) | |

| | | | |
|---|---|---|---|
| PKMYT1 | Protein Kinase, Membrane Associated Tyrosine/Threonine 1 | (368) | (369) |
| FANCD2 | FA Complementation Group D2 | (370) | |
| RBL1 | RB Transcriptional Corepressor Like 1 | (371) | |
| GLUL | Glutamate-Ammonia Ligase | (372) | (373) |
| RBMS3 | RNA Binding Motif Single Stranded Interacting Protein 3 | (374-377) | |
| TPM1 | Tropomyosin 1 | (378-382) | |
| CCDC8 | Coiled-Coil Domain Containing 8 | (383) | |
| MSH6 | MutS Homolog 6 | (312, 384) | |
| MEIS1 | Meis Homeobox 1 | (385-388) | |
| NSMAF | Neutral Sphingomyelinase Activation Associated Factor | (389-393) | |
| MYH10 | Myosin Heavy Chain 10 | (394) | |
| RBBP7 | RB Binding Protein 7, Chromatin Remodeling Factor | (395-397) | (398, 399) |
| TCF4 | Transcription Factor 4 | (400) | (401, 402) |
| TAF9B | TATA-Box Binding Protein Associated Factor 9b | (403) | (404) |
| HTR2B | 5-Hydroxytryptamine Receptor 2B | (405-407) | (408) |
| CDKN2C | Cyclin Dependent Kinase Inhibitor 2C | (409-412) | |
| LRIG3 | Leucine Rich Repeats And Immunoglobulin Like Domains 3 | (413-415) | |
| EXO1 | Exonuclease 1 | (416-418) | |
| PATZ1 | POZ/BTB And AT Hook Containing Zinc Finger 1 | (419-422) | |
| MXRA8 | Matrix Remodeling Associated 8 | (423, 424) | |
| RUNX1T1 | RUNX1 Partner Transcriptional Co-Repressor 1 | (425-427) | |
| ARHGAP11A | Rho GTPase Activating Protein 11A | (428) | |
| SLFN11 | Schlafen Family Member 11 | (429-433) | (434) |

61. Li B, Zheng Z, Wei Y, Wang M, Peng J, Kang T, et al. Therapeutic effects of neuregulin-1 in diabetic cardiomyopathy rats. Cardiovascular Diabetology. 2011;10(1):69.
62. Jie B, Zhang X, Wu X, Xin Y, Liu Y, Guo Y. Neuregulin-1 suppresses cardiomyocyte apoptosis by activating PI3K/Akt and inhibiting mitochondrial permeability transition pore. Molecular and Cellular Biochemistry. 2012;370(1):35-43.
63. Mòdol-Caballero G, Santos D, Navarro X, Herrando-Grabulosa M. Neuregulin 1 Reduces Motoneuron Cell Death and Promotes Neurite Growth in an in Vitro Model of Motoneuron Degeneration. Frontiers in Cellular Neuroscience. 2018;11.
64. Fock V, Plessl K, Draxler P, Otti GR, Fiala C, Knöfler M, et al. Neuregulin-1-mediated ErbB2–ErbB3 signalling protects human trophoblasts against apoptosis to preserve differentiation. Journal of Cell Science. 2015;128(23):4306-16.
65. Ohkoshi E, Umemura N. Induced overexpression of CD44 associated with resistance to apoptosis on DNA damage response in human head and neck squamous cell carcinoma cells. Int J Oncol. 2017;50(2):387-95.
66. Lakshman M, Subramaniam V, Rubenthiran U, Jothy S. CD44 promotes resistance to apoptosis in human colon cancer cells. Exp Mol Pathol. 2004;77(1):18-25.
67. Rottmann S, Speckgens S, Lüscher-Firzlaff J, Lüscher B. Inhibition of apoptosis by MAD1 is mediated by repression of the PTEN tumor suppressor gene. Faseb j. 2008;22(4):1124-34.
68. Foley KP, McArthur GA, Quéva C, Hurlin PJ, Soriano P, Eisenman RN. Targeted disruption of the MYC antagonist MAD1 inhibits cell cycle exit during granulocyte differentiation. Embo j. 1998;17(3):774-85.
69. Gao JX, Li Y, Wang SN, Chen XC, Lin LL, Zhang H. Overexpression of microRNA-183 promotes apoptosis of substantia nigra neurons via the inhibition of OSMR in a mouse model of Parkinson's disease. Int J Mol Med. 2019;43(1):209-20.
70. Sharanek A, Burban A, Laaper M, Heckel E, Joyal J-S, Soleimani VD, et al. OSMR controls glioma stem cell respiration and confers resistance of glioblastoma to ionizing radiation. Nature Communications. 2020;11(1):4116.
71. Busser B, Sancey L, Josserand V, Niang C, Favrot MC, Coll JL, et al. Amphiregulin promotes BAX inhibition and resistance to gefitinib in non-small-cell lung cancers. Mol Ther. 2010;18(3):528-35.
72. Kim D, Dai J, Fai LY, Yao H, Son YO, Wang L, et al. Constitutive activation of epidermal growth factor receptor promotes tumorigenesis of Cr(VI)-transformed cells through decreased reactive oxygen species and apoptosis resistance development. J Biol Chem. 2015;290(4):2213-24.
73. Ogata-Suetsugu S, Yanagihara T, Hamada N, Ikeda-Harada C, Yokoyama T, Suzuki K, et al. Amphiregulin suppresses epithelial cell apoptosis in lipopolysaccharide-induced lung injury in mice. Biochem Biophys Res Commun. 2017;484(2):422-8.
74. Liu K, Lin D, Ouyang Y, Pang L, Guo X, Wang S, et al. Amphiregulin impairs apoptosis-stimulating protein 2 of p53 overexpression-induced apoptosis in hepatoma cells. Tumour Biol. 2017;39(3):1010428317695026.
75. Wu F, Tian F, Zeng W, Liu X, Fan J, Lin Y, et al. Role of peroxiredoxin2 downregulation in recurrent miscarriage through regulation of trophoblast proliferation and apoptosis. Cell Death Dis. 2017;8(6):e2908.
76. Li H, Yang H, Wang D, Zhang L, Ma T. Peroxiredoxin2 (Prdx2) Reduces Oxidative Stress and Apoptosis of Myocardial Cells Induced by Acute Myocardial Infarction by Inhibiting the TLR4/Nuclear Factor kappa B (NF-κB) Signaling Pathway. Med Sci Monit. 2020;26:e926281.
77. Yang S, Luo A, Hao X, Lai Z, Ding T, Ma X, et al. Peroxiredoxin 2 inhibits granulosa cell apoptosis during follicle atresia through the NFKB pathway in mice. Biol Reprod. 2011;84(6):1182-9.
78. Zhou S, Han Q, Wang R, Li X, Wang Q, Wang H, et al. PRDX2 protects hepatocellular carcinoma SMMC-7721 cells from oxidative stress. Oncol Lett. 2016;12(3):2217-21.

219. Baptiste-Okoh N, Barsotti Anthony M, Prives C. A role for caspase 2 and PIDD in the process of p53-mediated apoptosis. Proceedings of the National Academy of Sciences. 2008;105(6):1937-42.
220. Vigneswara V, Akpan N, Berry M, Logan A, Troy CM, Ahmed Z. Combined suppression of CASP2 and CASP6 protects retinal ganglion cells from apoptosis and promotes axon regeneration through CNTF-mediated JAK/STAT signalling. Brain. 2014;137(6):1656-75.
221. Tiwari M, Sharma LK, Vanegas D, Callaway DA, Bai Y, Lechleiter JD, et al. A nonapoptotic role for CASP2/caspase 2. Autophagy. 2014;10(6):1054-70.
222. LeBlanc A, Liu H, Goodyer C, Bergeron C, Hammond J. Caspase-6 role in apoptosis of human neurons, amyloidogenesis, and Alzheimer's disease. J Biol Chem. 1999;274(33):23426-36.
223. Cowling V, Downward J. Caspase-6 is the direct activator of caspase-8 in the cytochrome c-induced apoptosis pathway: absolute requirement for removal of caspase-6 prodomain. Cell Death Differ. 2002;9(10):1046-56.
224. Luo T, Yuan Y, Yu Q, Liu G, Long M, Zhang K, et al. PARP-1 overexpression contributes to Cadmium-induced death in rat proximal tubular cells via parthanatos and the MAPK signalling pathway. Scientific Reports. 2017;7(1):4331.
225. Spindel ON, World C, Berk BC. Abstract 13533: Thioredoxin Interacting Protein Prevents Endothelial Cell Apoptosis - Critical Mediator of VEGF Receptor 2 Activation via Poly-ADP Ribose Polymerase 1. Circulation. 2011;124(suppl_21):A13533-A.
226. Sahaboglu A, Tanimoto N, Kaur J, Sancho-Pelluz J, Huber G, Fahl E, et al. PARP1 Gene Knock-Out Increases Resistance to Retinal Degeneration without Affecting Retinal Function. PLOS ONE. 2010;5(11):e15495.
227. Chiarugi A, Moskowitz MA. PARP-1--a Perpetrator of Apoptotic Cell Death? Science. 2002;297(5579):200-1.
228. Los M, Mozoluk M, Ferrari D, Stepczynska A, Stroh C, Renz A, et al. Activation and Caspase-mediated Inhibition of PARP: A Molecular Switch between Fibroblast Necrosis and Apoptosis in Death Receptor Signaling. Molecular Biology of the Cell. 2002;13(3):978-88.
229. Zhou L, Cai X, Han X, Xu N, Chang DC. CDK1 switches mitotic arrest to apoptosis by phosphorylating Bcl-2/Bax family proteins during treatment with microtubule interfering agents. Cell Biol Int. 2014;38(6):737-46.
230. Furukawa Y, Iwase S, Terui Y, Kikuchi J, Sakai T, Nakamura M, et al. Transcriptional activation of the cdc2 gene is associated with Fas-induced apoptosis of human hematopoietic cells. J Biol Chem. 1996;271(45):28469-77.
231. Choi KS, Eom YW, Kang Y, Ha MJ, Rhee H, Yoon J-W, et al. Cdc2 and Cdk2 Kinase Activated by Transforming Growth Factor-β1 Trigger Apoptosis through the Phosphorylation of Retinoblastoma Protein in FaO Hepatoma Cells *. Journal of Biological Chemistry. 1999;274(45):31775-83.
232. Gu L, Zheng H, Murray SA, Ying H, Jim Xiao ZX. Deregulation of Cdc2 kinase induces caspase-3 activation and apoptosis. Biochem Biophys Res Commun. 2003;302(2):384-91.
233. Wang S, Hasham MG, Isordia-Salas I, Tsygankov AY, Colman RW, Guo Y-L. Upregulation of Cdc2 and cyclin A during apoptosis of endothelial cells induced by cleaved high-molecular-weight kininogen. American Journal of Physiology-Heart and Circulatory Physiology. 2003;284(6):H1917-H23.
234. Wu J, Feng Y, Xie D, Li X, Xiao W, Tao D, et al. Unscheduled CDK1 activity in G1 phase of the cell cycle triggers apoptosis in X-irradiated lymphocytic leukemia cells. Cell Mol Life Sci. 2006;63(21):2538-45.
235. Matthess Y, Raab M, Sanhaji M, Lavrik IN, Strebhardt K. Cdk1/cyclin B1 controls Fas-mediated apoptosis by regulating caspase-8 activity. Mol Cell Biol. 2010;30(24):5726-40.
236. Wang J, He N, Wang R, Tian T, Han F, Zhong C, et al. Analysis of TET2 and EZH2 gene functions in chromosome instability in acute myeloid leukemia. Scientific Reports. 2020;10(1):2706.
