## Supplementary figures and images for "Connectivity Map and perturbation-based sensitivity analysis identifies MEK inhibitors as senolytics in human lung fibroblasts"

### Fig S1

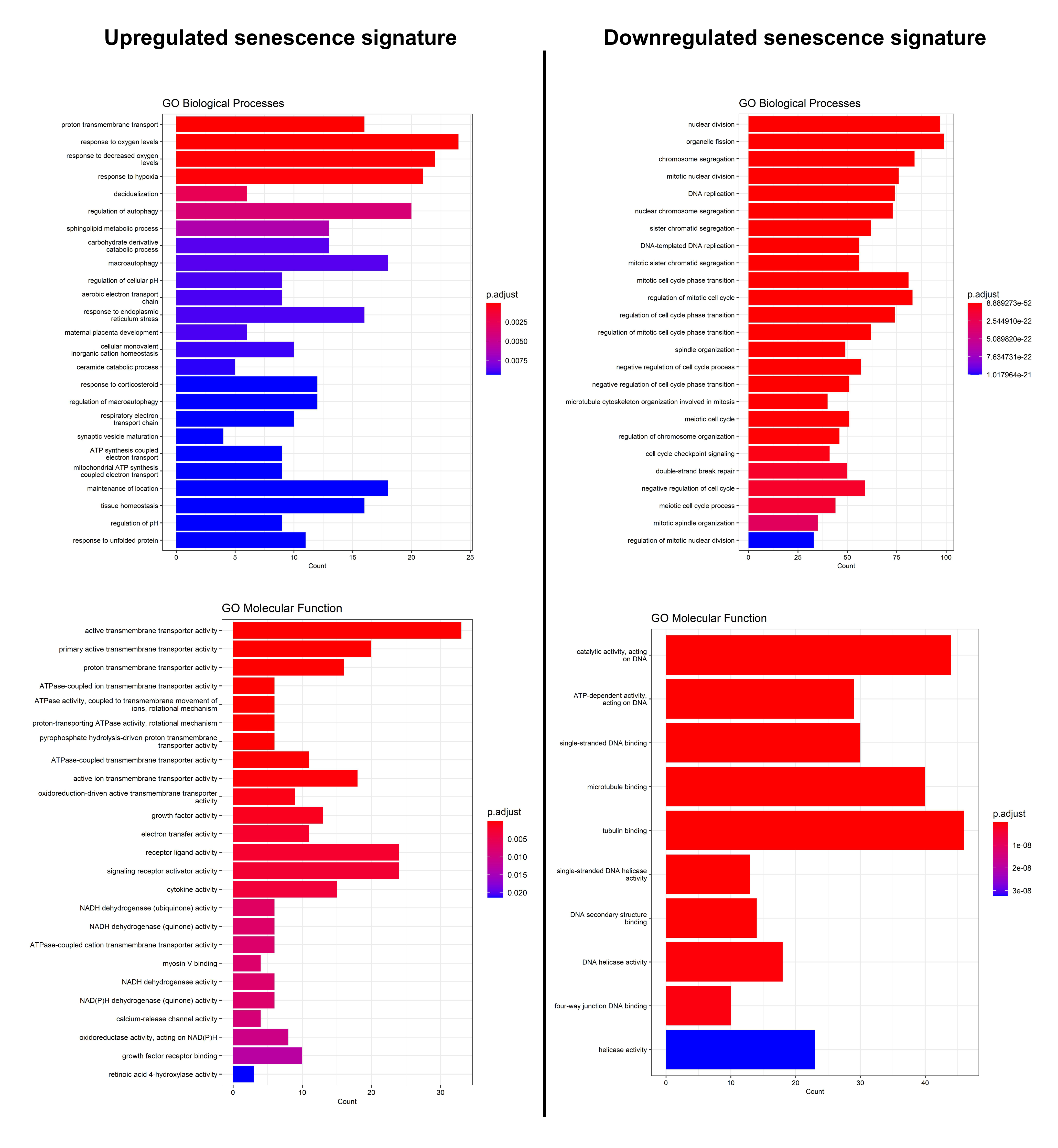
